# Extreme distributions in the preconfigured developing brain

**DOI:** 10.1101/2023.11.13.566810

**Authors:** Mattia Chini, Marilena Hnida, Johanna K. Kostka, Yu-Nan Chen, Ileana L. Hanganu-Opatz

**Author notes:** Corresponding author: Mattia Chini.

## Abstract

In the adult brain, structural and functional parameters, such as synaptic sizes and neuronal firing rates, follow right-skewed and heavy-tailed distributions. While this organization is thought of having significant implications, its development is still largely unknown. Here, we address this knowledge gap by investigating a large-scale dataset recorded from the prefrontal cortex (PFC) and the olfactory bulb of mice aged 4-60 postnatal days. We show that firing rates and pairwise correlations have a largely stable distribution shape over age, and that neural activity displays a small-world architecture. Moreover, early brain activity displays an oligarchical organization, i.e., neurons with high firing rates are likely to have hub-like properties. Leveraging neural network modeling, we show that analogously extremely distributed synaptic parameters are necessary to recapitulate the experimental data. Thus, functional and structural parameters in the developing brain are already extremely distributed, suggesting that this organization is preconfigured and not experience-dependent.

## Introduction

In the adult brain, many structural and functional parameters follow right-skewed distributions with heavy right tails such as the log-normal^1,2^, gamma^3^, or power law^4,5^ distribution. A non-exhaustive list of parameters that is characterized by such distributions includes: size and number of synapses^6–9^, size of post-synaptic currents^10– 12^, diameter of axons^13^, density of neurons^14^, *in vivo* single-unit firing rate^15–17^, spike transmission probability^10,17^, pairwise correlations among spike trains^18^, and power of the LFP^19,20^, EEG^21–23^, MEG^22–24^ and BOLD^25^ signals.

If a parameter follows such distributions, it is implied that a large proportion of the data displays small values that fall well below the mean. Conversely, extremely large values are more commonly observed than if it would follow a narrower distribution such as the normal (or Gaussian), as it is often implicitly assumed^1,2^. Thus, these distributions are characterized by high levels of inequality. We define such distributions as being “extreme”^26^. It has been suggested that extreme distributions of neural parameters might have several useful properties^1,2,10,27^. For instance, the log-normal distribution of synaptic sizes might promote the formation and propagation of neuronal sequences^10,28^, while at the same time optimizing storage capacity^27^. Moreover, the extreme distribution of firing rates might result in an environment with an optimal balance between a large amount of “specialist” neurons complemented by few “generalists”. The first ones would only fire upon receiving a highly specific constellation of presynaptic inputs, and thereby have a spiking activity with unique and distinctive “interpretation” for postsynaptic partners. Conversely, “generalist” neurons would require less specific presynaptic inputs to generate a spike. This could allow such neurons to generalize over similar sensory stimuli and might therefore represent the brain’s “best guess”^1,2^. Overall, this organization is thought of producing a system that allows for concomitant specialization and flexibility, while limiting the number of energy-demanding high firing rate neurons^10^.

While it is widely accepted that the adult brain is an extreme environment, how this unfolds throughout development is still unclear. Two main competing hypotheses have been put forward. The so-called “blank slate model” posits that the developing brain is a “*tabula rasa*”, a blank slate that lacks a refined structure. A corollary of this view is that structural and functional parameters in the developing brain should follow a narrow, thin-tailed distribution. Only upon developmental learning would the brain structure gradually become skewed, heavy-tailed, and inequal, as reported for the adult brain. While this view is not often openly advocated for, it is often implied^2,29,30^. The competing theory is the “preconfigured brain hypothesis”, that proposes that the developing brain is pre-packaged with non-random structure and already displays extreme distributions of structural and functional parameters. Consequently, this hypothesis implies that developmental learning should not fundamentally transform the brain architecture. Rather, learning is viewed as a matching process that associates pre-existing structure, that is initially devoid of meaning, with a behavioral output or a sensory sensation^2,31–34^.

To tease apart these two hypotheses, we investigated the *in vivo* development of brain activity in a large-scale dataset from unanesthetized postnatal day (P) 4-60 mice (n=302 mice). The youngest mice in the dataset belong to a very immature developmental phase in which, even though neurons in the sensory systems are active, they mostly do not represent sensory information. An exception to this developmental dynamic is the olfactory system. From birth on, rodents rely on olfaction for survival, as they actively employ olfactory cues to locate the dam’s nipple^35,36^. Thus, in the present study, to control for the potential effect of developmental learning, we investigated the early developing olfactory bulb (OB) and the prefrontal cortex (PFC), the cortical region with the most protracted development^37^. We show that, in both brain regions, the skewness, tailedness and inequality of the distributions of *in vivo* single unit activity (SUA) firing rates and pairwise interactions among them do not increase throughout development. Along the same lines, already midway through the first postnatal week, the PFC displays a non-random small-world topography. Moreover, in both brain regions, neurons in the right tail of the firing rate distribution are overwhelmingly more likely to also be in the right tail of the pairwise interaction distribution, and to exhibit hub-like properties. We refer to this organization as oligarchical, and we show that it gradually wanes throughout adulthood, concomitantly with a maturation of excitation-inhibition balance.

## Results

### SUA firing rate exponentially increases over age, while spiking activity decorrelates

To assess the *in vivo* developmental firing dynamics, we analyzed a large-scale dataset of SUA (n=278 mice, 14357 units and 654335 spike train pairs) recorded with Neuropixels as well as single- and multi-shank Neuronexus silicon probes from the PFC and the ventral OB of non-anesthetized P4-P12 mice (n=278 mice) (Figure 1A-C, Supp. Figure 1A-C). Part of the data has been used in previously published studies^38,39^. We calculated the firing rate (first-order SUA statistics) and the spike-time tiling coefficient (STTC, second-order SUA statistics, calculated at a 10 ms timescale), a measure of pairwise correlation among spike trains that is not biased by firing rate^40^. In line with previous reports, using multi-variate linear regression, we found that firing rate exponentially increased with age in both brain regions (age effect=0.11, CI [0.04; 0.18], p=0.004, generalized linear mixed-effect model) (Figure 1D) and that the OB displayed a higher firing rate than the PFC (firing rate at P8=0.28 and 0.04 Hz, CI [0.21; 0.37] and [0.03; 0.06] for OB and PFC, respectively, p<10^-16^) (Figure 1D).

**Figure 1.**
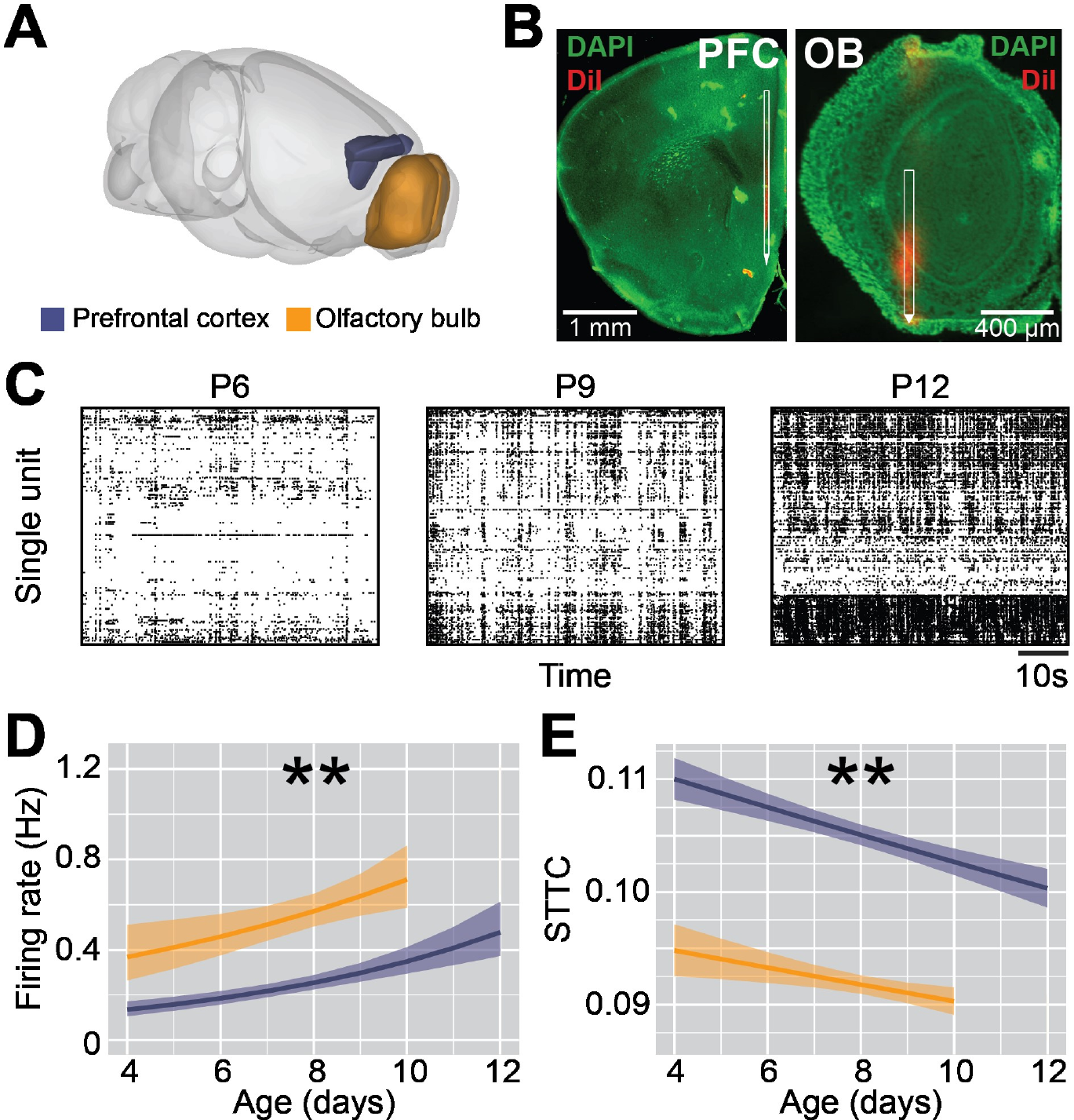
SUA firing statistics in the mouse PFC and OB across the first two postnatal weeks. (**A**) Schematic representation^41^ of extracellular recordings in the PFC and OB of P4-12 mice. (**B**) Digital reconstruction of the position of a DiI-labeled recording electrode in the PFC (left) and OB (right) of a Nissl-stained coronal section (green) of a P9 and P10 mouse, respectively. (**C**) Representative raster plot of 1 minute of SUA activity recorded in the PFC of a P6 (left), P9 (middle) and P12 (right) mouse. (**D**) Line plot displaying the SUA firing rate of P4-12 mice (n=278 mice and 14357 single units). Color codes for brain region. (**E**) Same as (**D**) for STTC (n=278 mice and 654335 spike train pairs). In (**D)** and (**E**) data is presented as mean and 95% confidence interval. Asterisks in (**D**), and (**E**) indicate significant effect of age. **p<0.01, generalized linear mixed-effect models.

Concomitantly to the increase of firing rate, and consistently with previously published data^38,42,43^, the STTC exponentially decreased with age (age effect=-0.008, CI [-0.014; −0.003], p=0.003, linear mixed-effect model), indicating a developmental decorrelation of spiking activity (Figure 1E). When evaluating this effect on individual brain regions, we found that the OB displayed lower STTC values than the PFC (STTC value at P8=0.0041 and 0.0056 Hz, CI [0.0040; 0.0042] and [0.0055; 0.0057] for OB and PFC, respectively, p<10^-16^) (Figure 1E).

This data indicates that, in both brain regions, the firing rates exponentially increase across the first two postnatal weeks while, at the same time, spiking activity decorrelates. Consistent with the early maturation of the OB and its high density of INs^36^, the OB display higher firing rates and lower STTC values than the PFC.

**Supplementary Figure 1.**
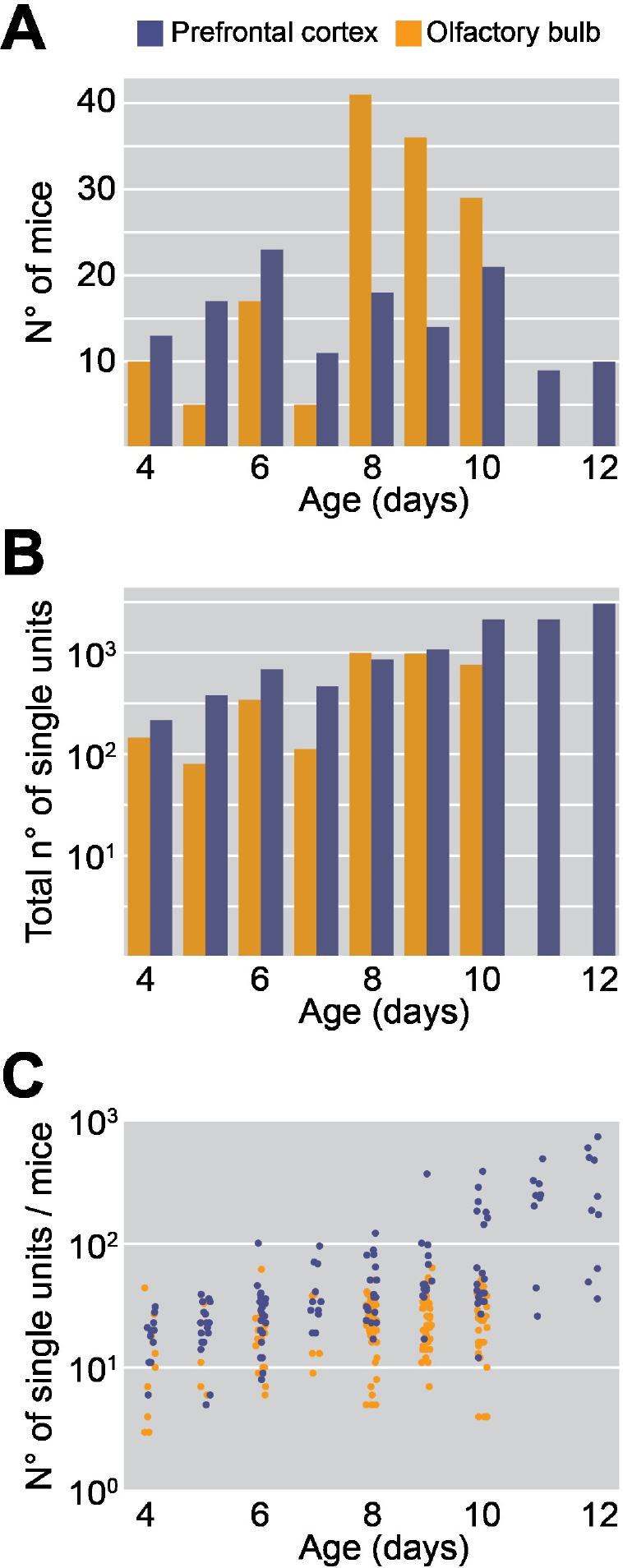
Descriptive statistics of the experimental dataset. (**A**) Bar plot displaying the number of recorded mice across age and brain region (n=278 mice). Color codes for brain region. (**B**) Same as A for recorded single units (n=14357 single units). (**C**) Scatter plot displaying the number of recorded units per individual mice over age and brain regions (n=278 mice). In (**C**) dots indicate individual mice.

### The skewness, tailedness and inequality of the SUA firing statistics do not increase over age

A corollary of the “blank slate model” is that the distributions of functional parameters should become more extreme over development. To quantify the distribution shape for the first and second-order SUA statistics, we computed their skewness, kurtosis and Gini coefficient (Figure 2A-C). Skewness and kurtosis are the 3^rd^ and 4^th^ central moments of a variable, and measure its asymmetry and tailedness, respectively. Negative values of skewness indicate a left-skewed distribution, positive values a right-skewed distribution, and null values a symmetrical distribution (Figure 2A). Kurtosis only takes positive values. Normal distributions have a kurtosis value of 3, and are referred to as mesokurtic distributions. Distributions with a kurtosis below 3 are platykurtic (thin-tailed), whereas distribution with a kurtosis above 3 are leptokurtic (heavy-tailed) (Figure 2B). The Gini coefficient measures the inequality of a distribution^44,45^ and takes values ranging between 0 (representing total equality) and 1 (representing total inequality). It is calculated as the area between the line of equality and the Lorenz curve divided by the total area under the line of equality (Figure 2C). To increase the estimation accuracy of these three parameters, for this analysis we only considered mice with at least 20 simultaneously recorded single units (n=195/278 mice).

**Figure 2.**
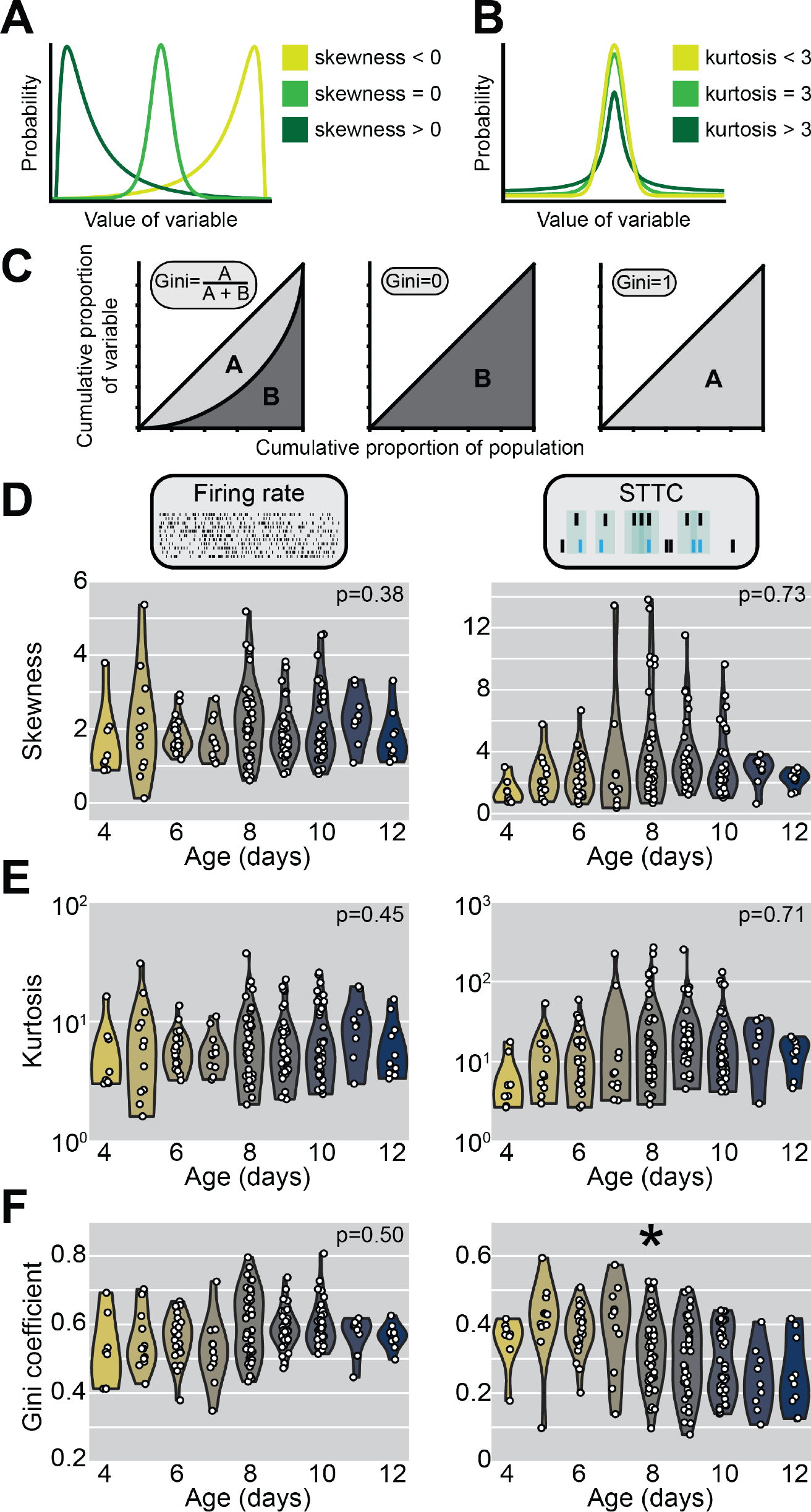
Skewness, kurtosis and Gini coefficient of firing rate and STTC over the first two postnatal weeks. (**A**) Schematic representation of 3 distributions with different skewness values. (**B**) Same as (**A**) for kurtosis. (**C**) Lorenz curves for 3 distributions with different Gini coefficient values and schematic representation of how the Gini coefficient is calculated. (**D**) Violin plot displaying the skewness of firing rate (left) and STTC (right) of P4-12 mice (n=195 mice). Color codes for age with 1-day increments. (**E-F**) Same as (**D**) for kurtosis and Gini coefficient. In (**E-F**) white dots indicate individual mice. In (**E-F**) the shaded area represents the probability density of the variable. Asterisks in (**F**) indicate a significant effect of age. **p<0.01, linear models.

Contrary to the prediction of the “blank slate model”, neither the skewness nor the kurtosis of firing rates in the PFC and OB (age coefficient=-0.06 and −0.02, CI [-0.19; 0.07] and [-0.05; 0.02], p=0.39 and p=0.45, for skewness and kurtosis, respectively, linear model) and STTC (age coefficient=0.06 and 0.01, CI [-0.27; 0.39] and [-0.05; 0.07], p=0.73 and p=0.71, for skewness and kurtosis, respectively, linear model) exhibited an age-dependent trend (Figure 2D-E). Pooling together firing rate and STTC distributions, 100% of the distributions were right-skewed (390/390, skewness > 0), and 94% were leptokurtic or heavy-tailed (366/390, kurtosis > 3). Along the same lines, the Gini coefficient of firing rate did not significantly change over the first two postnatal weeks (age coefficient=-0.004, CI [-0.014; 0.007], p=0.50, linear model), whereas the Gini coefficient of STTC even slightly decreased over age (age coefficient=-0.02, CI [-0.03; −0.002], p=0.02, linear model) (Figure 2F). This dynamic was consistent across brain regions (Supp. Figure 2A-C) and robust to changes in the minimum number of single units used as cutoff for analysis (Supp. Figure 2D). Even analyzing the distributions of firing rate and STTC after pooling together units recorded in the same brain region and in mice of the same age did not lead to age-dependency for any of the evaluated parameters (Supp. Figure 3).

**Figure 3.**
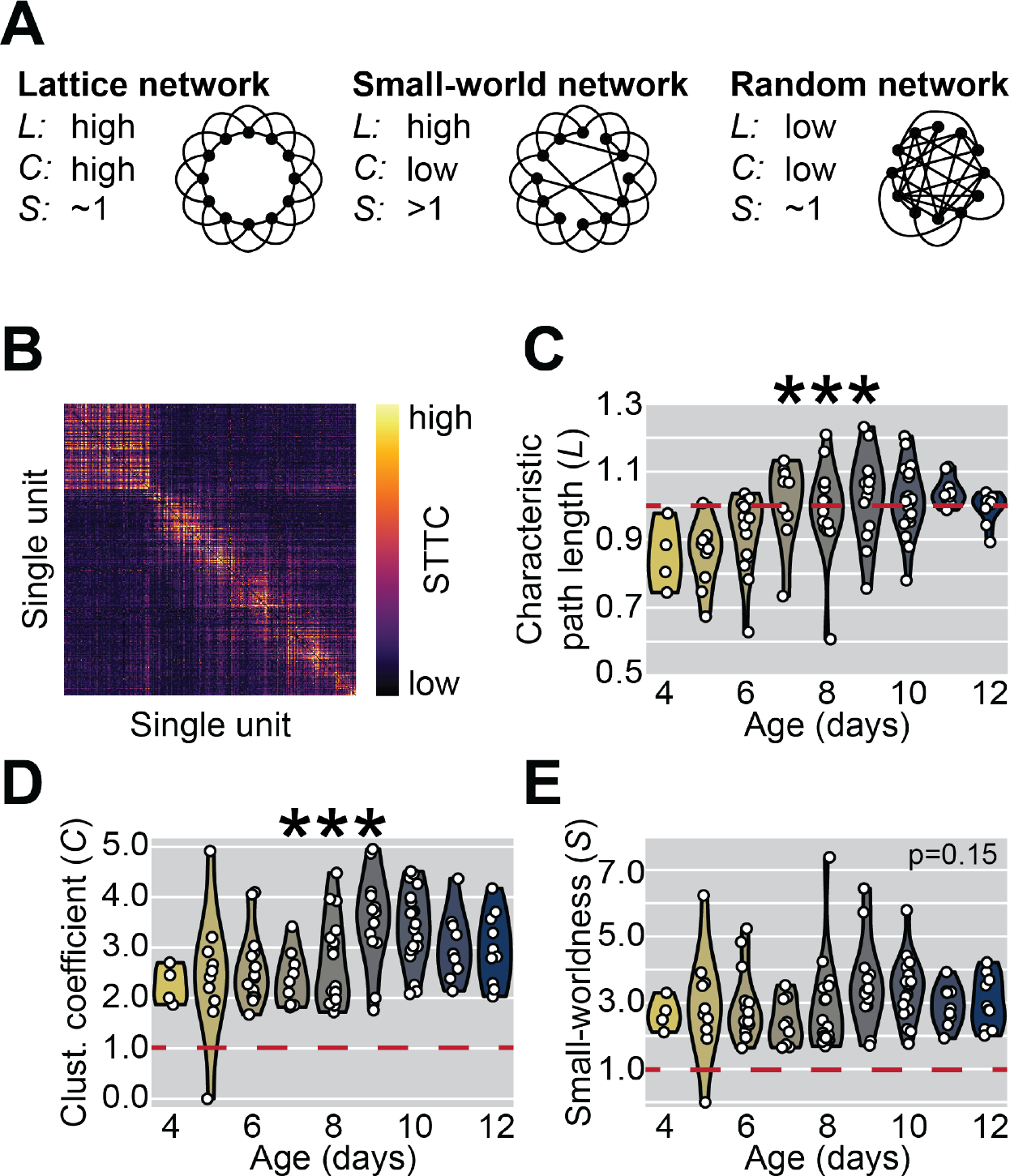
Complex network analysis in the PFC across the first two postnatal weeks. (**A**) Schematic representation of a lattice, small-world and random network and their respective values of characteristic path length (L), clustering coefficient (C) and small-worldness (S). (**B**) Weighted adjacency displaying a representative STTC matrix of a P10 mouse. Color codes for STTC value. Units are sorted by recording depth. (**C**) Violin plot displaying the characteristic path length as a function of age (n=108 mice). Color codes for age with 1-day increments. (**D-E**) Same as (**C**) for clustering coefficient and small-worldness. In (**C-E**) white dots indicate individual mice and shaded area represents the probability density of the variable. Asterisks in (**C-D**) indicate a significant effect of age. ***p<0.001, linear models.

**Supplementary Figure 2.**
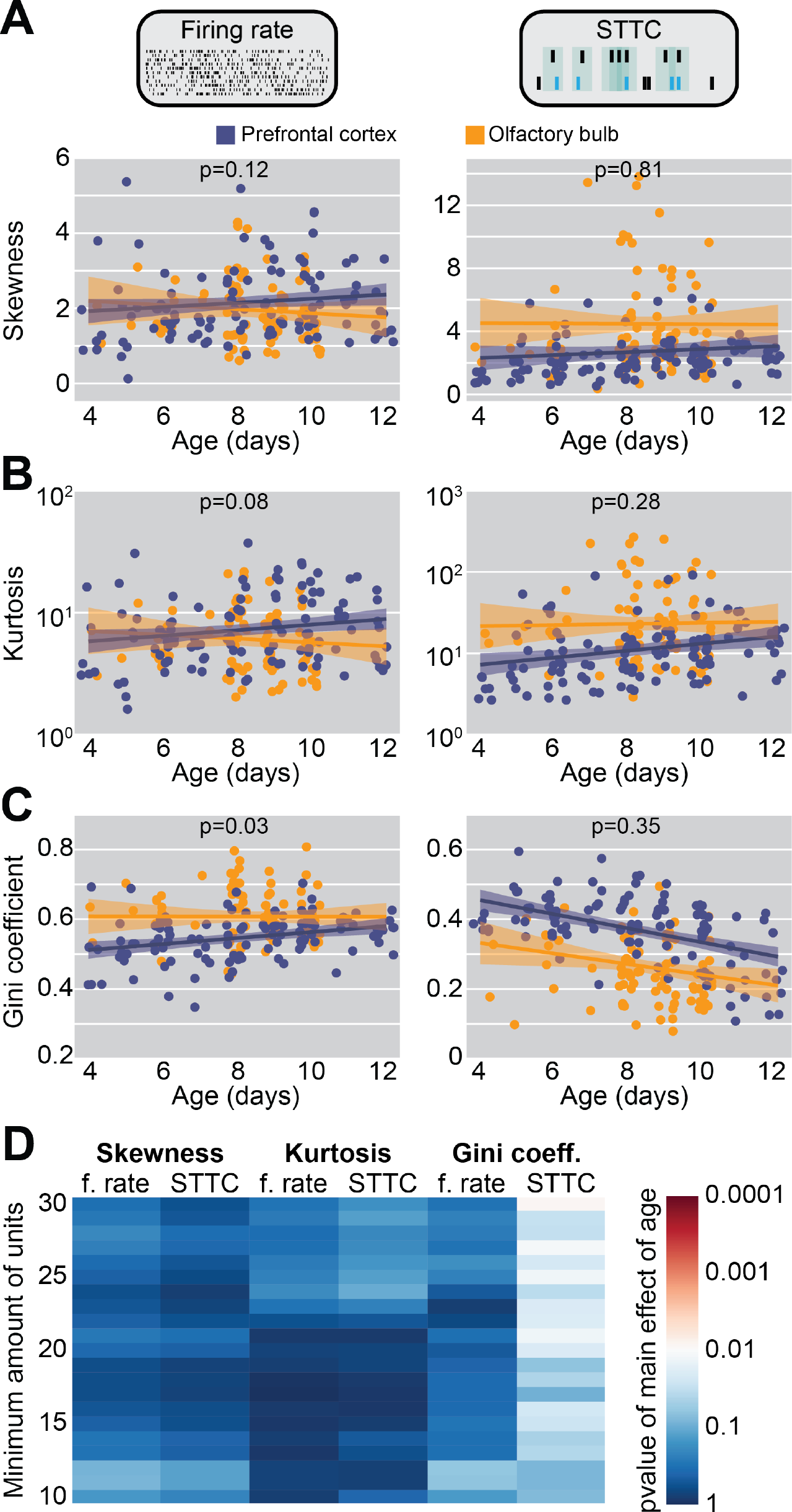
Skewness, kurtosis and Gini coefficient of firing rate and STTC across brain regions in the first two postnatal weeks. (**A**) Scatter and line plot displaying the skewness of firing rate (left) and STTC (right) of P4-12 mice (n=238 mice). Color codes for brain region. (**B-C**) Same as (A) for kurtosis and Gini coefficient. (**D**) Heatmap displaying the p-value for the main effect of age as a function of the minimum number of units used as a cutoff to be included in the analysis. In (**A-C**) colored dots indicate individual mice, and data is presented as mean and 95% confidence interval. P-values refer to the interaction between age and brain region.

**Supplementary Figure 3.**
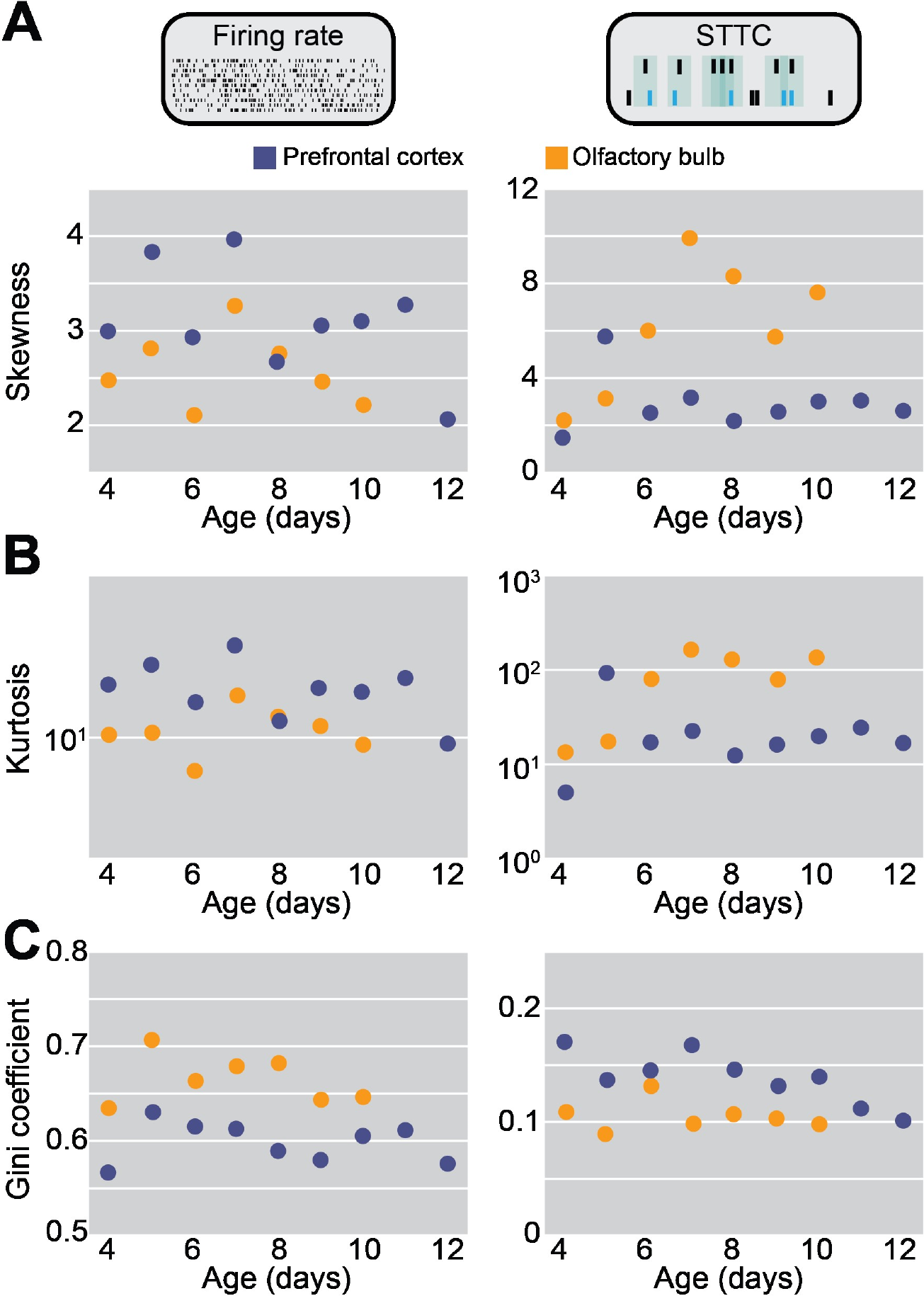
Skewness, kurtosis and Gini coefficient of firing rate and STTC pooled across mice over the first two postnatal weeks. (**A**) Scatter plot displaying the skewness of firing rate (left) and STTC (right) of P4-12 mice. Color codes for brain region. (**B-C**) Same as (**A**) for kurtosis and Gini coefficient. In (**A-C**) dots indicate a single parameter estimation on data pooled across mice of the same age recorded from the same brain region.

This data indicates that, irrespective of brain region, the skewness, tailedness and inequality of first- and second-order SUA statistics do not increase with age, as would be predicted by the “blank slate model”. On the contrary, the only parameter to exhibit a significant age-dependence is the Gini coefficient of STTC, that modestly decreases with age. Further, almost the totality of these distributions is right-skewed and heavy-tailed. The fact that these results are largely brain region-independent indicates that the different developmental speed of the early maturating OB and the late maturing PFC does not affect these dynamics. Considering the large size of the investigated datasets, it is also unlikely that we were not able to detect the presence of developmental trends due to lack of statistical power.

Lastly, solely skewness and kurtosis of both firing rate and STTC robustly correlated with each other (r2=0.9 and 0.88, respectively). Despite similar developmental trajectories, the other pairwise parameters combination were poorly predictive of each other (median r2=0.03) (Supp. Figure 4), indicating that they quantify distinct distributions that are largely independent of each other and thus, not redundant.

**Supplementary Figure 4.**
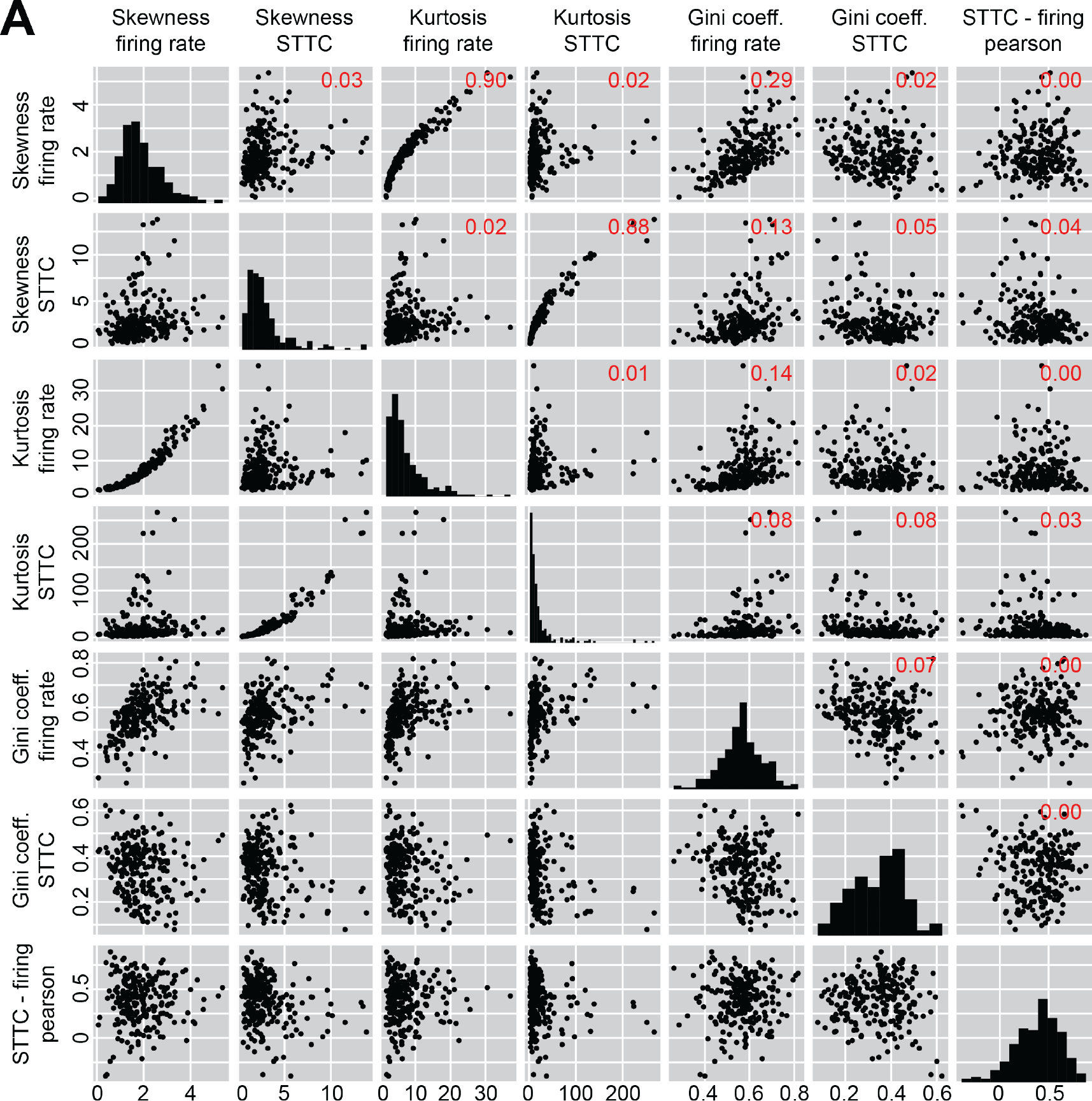
Distribution and pairwise correlations of skewness, kurtosis and Gini coefficient of firing rate and STTC. (**A**) Pair plot of the 7 parameters used to describe the shape of the firing rate and STTC distributions. The plots on the main diagonal displays histograms of the distributions of each individual parameter across the entire dataset (n=238 mice). The off-diagonal plots display the pairwise correlations among the 7 parameters. The numbers in red indicate the Pearson r2 for that specific pairwise parameter combination. Black dots represent individual mice.

**Figure 4.**
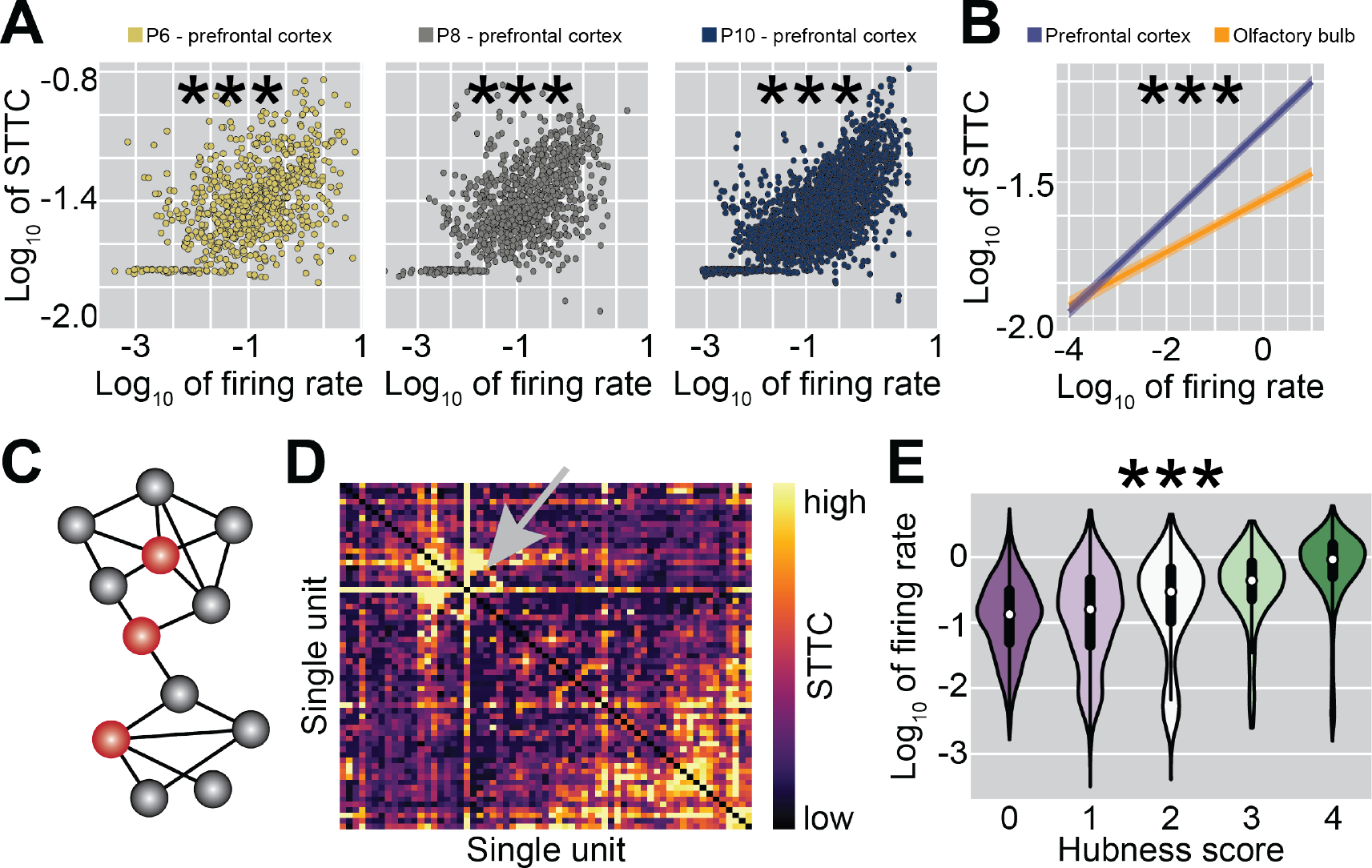
Firing rate correlates with average STTC and hubness score. (**A**) Scatter plot displaying the log-transformed average STTC of prefrontal neurons as a function of their log-transformed firing rate in all recorded P6 (left), P8 (center) and P10 (right) mice. Color codes for age. (**B**) Line plot displaying the log-transformed average STTC as a function of log-transformed firing rate across brain regions (n=259 mice and 14043 single units). Color codes for brain region. (**C**) Schematic representation of a network’s graph in which regional and global hubs are colored in red. (**D**) Weighted adjacency STTC matrix recorded from the PL of a P10 mouse. The gray arrow indicates a neuron with high hubness score. Color codes for STTC value. (**E**) Violin plot displaying the log-transformed firing rate of a neuron as a function of its hubness score. Color codes for hubness score. In (**A**) colored dots indicate individual neurons. In (**E**) data is presented as median, 25^th^, 75^th^ percentile, and interquartile range, and the shaded area represents the probability density of the variable. Asterisks in (**A-B**) indicate a significant effect of age, in (**E**) of hubness score. ***p<0.001, linear mixed-effect models.

### Complex network properties of the PFC do not vary over development

Even if the skewness, tailedness and inequality of firing rate and STTC are largely constant over age, we hypothesized that developmental changes might be detectable at a network level. To test this hypothesis, we resorted to complex network analysis^46^, and investigated the network topology of the developing PFC *in vivo* (Figure 3A), similarly to what previously done *in vitro*^47^. To minimize a potential estimation bias due to low number of single units, an inherent drawback of recordings in neonatal mice, we limited this analysis to mice having at least 20 single units (n=108/131 PFC recordings). We restricted the investigation to the PFC because the network analysis for OB was biased by the low number of units and the level of recurrent connectivity.

The network analysis was carried out on symmetric STTC matrices computed on individual mice, where each node corresponds to a single unit (Figure 3B). These matrices were thresholded and binarized by a shuffling procedure that swapped the identity of the neurons while preserving the population firing rate (see Materials and Methods for details). To evaluate the topology of the graphs, we computed their density and three main network properties: characteristic path length (L), clustering coefficient (C) and small-worldness (S) (Figure 3A). These last three parameters were normalized by dividing them with a corresponding null value extracted from random networks with the same density.

Similar to the parameters based on SUA statistics, the density of graphs did not significantly vary over age (age coefficient=-0.003, CI [-0.009; 0.002], p=0.23, linear model) (Supp. Figure 5A). While L values were similar to those computed on random networks (58/108=54% networks with normalized C > 1) (Figure 3C), all but one network had larger C values than the corresponding random network (107/108=99% networks with normalized L > 1) (Figure 3D). The graphs’ transitivity, a parameter that is closely related to the clustering coefficient, was analogously higher than in random networks (107/108=99% networks with normalized transitivity > 1) (Supp. Figure 5B).

**Figure 5.**
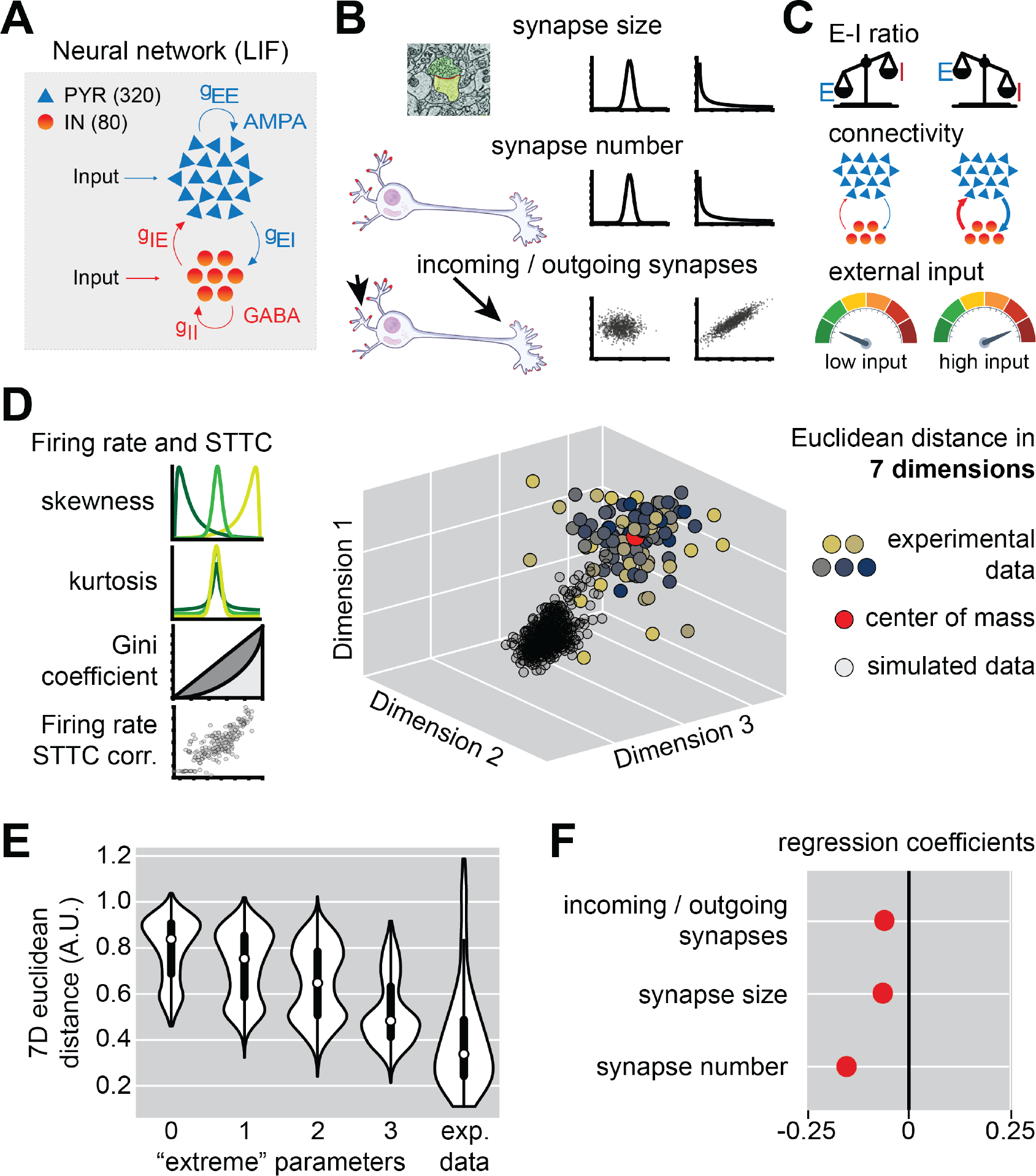
Spiking neural network modeling of the of the distribution shape of SUA statistics in the developing PFC. (**A**) Schematic representation of the neural network model. (**B-C**) Same as A for the 3 synaptic parameters (**B**) and the simulation parameters that were treated as random variables (**C**). (**D**) Schematic representation of the parameters that were used to evaluate the distribution shape of the experimental and simulated spiking data (left) and the approach that was used to evaluate the distance between experimental and simulated data (right). Note that, even though only 3 dimensions are shown in the scatter plot, the distance between simulated and experimental data was calculated in a 7-dimensional space. Color in the scatter plot codes for age. (**E**) Violin plot displaying the distance between simulated data and the center of mass of experimental data as a function of the number of synaptic parameters set in their “extreme” configuration. (**F**) Multivariate regression coefficients for the 3 synaptic parameters over the distance from the center of mass of experimental data. In (**E**) data is presented as median, 25^th^, 75^th^ percentile, and interquartile range, and the shaded area represents the probability density of the variable. In (**F**) regression coefficients are presented as mean and 95% confidence interval.

Low L and high C values are typical of so-called small-world networks, a category that includes many real world networks, including the adult brain^48–50^ (but see^51^). Accordingly, the small-worldness of the developing PFC was robustly higher than its corresponding null value (107/108=108=99% networks with normalized S > 1) (Figure 3E). While the normalized L and C increased over age (age coefficient=0.02 and 0.13, CI [0.01; 0.03] and [0.06; 0.21], p<10^-4^ and p<10^-4^, for normalized C and L, respectively, linear model) (Figure 3C-D), the normalized S did not vary with age (age coefficient=0.07, CI [-0.02; 0.16], p=0.15, linear model) (Figure 3E).

Thus, complex network analysis revealed that already during the first two postnatal weeks, the PFC displays a non-random and small-world network architecture that is similar to that described in the adult brain^48–50^ and does not evolve with age.

**Supplementary Figure 5.**
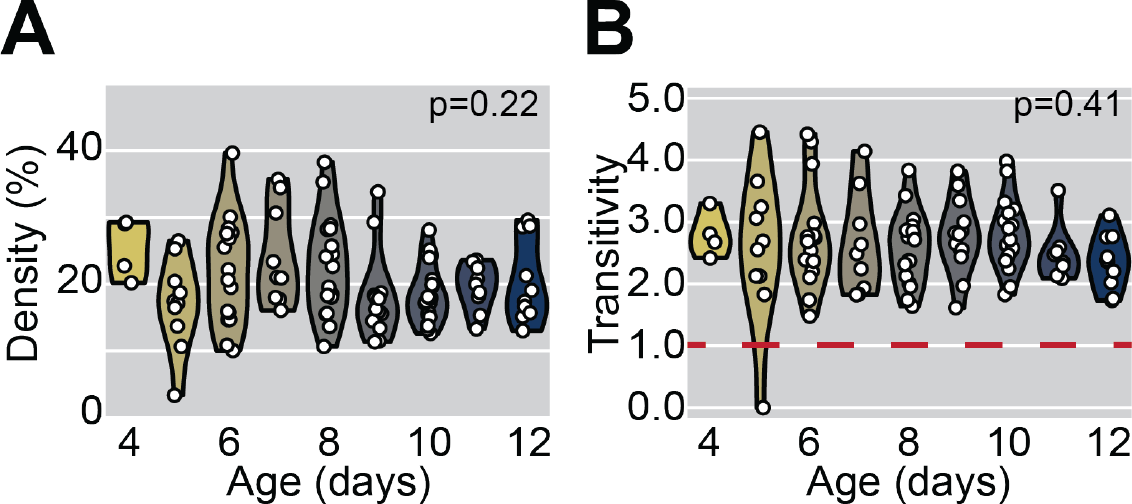
Graph density and transitivity of the PFC across the first two postnatal weeks. (**A**) Violin plot displaying the graph density of P4-12 mice (n=108 mice). Color codes for age with 1-day increments. (**B**) Same as (**A**) for transitivity. In (**A-B**) white dots indicate individual mice. In (**A-B**) the shaded area represents the probability density of the variable.

### The developing brain is in an oligarchical state

Next, we investigated whether the SUA statistics are not only extremely distributed, but also correlated to each other. For this, we used multivariate linear regression with age as a covariate, and evaluated the relationship between the log-transformed firing rate of a neuron and its log-transformed average STTC. We found that, irrespective of brain region, the two variables robustly correlated with each other (firing rate coefficient=0.08, CI [0.07; 0.08], p<10^-70^, linear mixed-effect model) (Figure 4A-B). This correlation between firing rate and average STTC was lower in the OB than in the PFC (firing rate and brain area interaction coefficient=0.06, CI [0.05; 0.06], p<10^-84^, linear mixed-effect model) (Figure 4B). Since the STTC lacks a firing rate bias^40^, this is indicative of a genuine correlation between the two variables. Consequently, neurons with high firing rates disproportionately contribute to network dynamics during development, a property reminiscent of hub neurons, as recently shown in the developing barrel cortex^52^. We define this state, in which extreme distributions are tightly correlated with each other, as being “oligarchical”.

To investigate whether the firing rate of a neuron also correlates to its network-level features, we used the prefrontal STTC matrices described in the paragraph above and extracted 4 different regional and global “hubness” metrics for each individual node: degree (i.e. total number of connections), strength (i.e. sum of connection weights), betweenness centrality (the fraction of shortest paths containing a given node) and closeness centrality (the reciprocal of the average shortest path for a given node) (Figure 4C-D). For each of these measures, we then assigned a score of 0 or 1 to each node, depending on whether it exceeded the 75^th^ percentile of that specific metric^47^. Summing the individual scores, we obtained a composite hubness score, with values ranging from 0 to 4. The majority of neurons (6580/10512=63%) had a hubness score of 0, whereas a smaller proportion (1471/10512=14%) of neurons received a hubness score of 4. As previously reported in 2D neuronal cultures^47^, the hubness score robustly correlated with firing rate (hubness score effect, p<10^-70^, linear mixed-effect model), with the median firing rate differing by almost an order of magnitude between neurons with hubness score 0 and 4 (Figure 5E).

These data indicate that the developing cortex is not only an environment characterized by stably extreme SUA statistics distributions, but also that it exhibits a peculiar oligarchical state, in which the firing rate of a neuron robustly correlates with the strength of its pairwise interactions and its network-level properties.

### Extreme synaptic distributions are necessary to recapitulate early PFC activity in a spiking neural network model

To investigate the mechanisms underlying the extreme distributions of SUA statistics, we hypothesized that analogously extremely distributed structural synaptic parameters might be necessary. To explore this proposition, we simulated ∼8000 spiking neural networks of interconnected conductance-based leaky integrate-and-fire (LIF) neurons. The simulated networks had no spatial structure, and consisted of 400 neurons, 80% of which were excitatory (PYRs) and 20% inhibitory (INs), in line with anatomical data for neocortical areas^53,54^. PYRs were simulated with outgoing excitatory AMPA synapses, while INs were equipped with outgoing inhibitory GABAergic synapses (Figure 5A). In these models, we investigated how the simulated firing statistics were impacted by three structural parameters: (i) whether the size of synapses followed a normal or a log-normal distribution, (ii) whether the number of synapses of individual neurons followed a normal or a log-normal distribution, and (iii) whether the number of dendritic and axonic (incoming and outgoing) synapses were correlated or uncorrelated with each other (Figure 5B). This resulted in a total of 2^3^=8 possible synaptic parameters combinations. To avoid choosing an arbitrary network architecture to test the effect of these three variables, a set of other parameters (average size of AMPA and GABA synapses, average connectivity, and the strength of the noisy input driving PYRs and INs) were randomly drawn from a range of biologically constrained values (see Materials and Methods for details) (Figure 5C). Given that the general neural network architecture is inspired by the neocortical anatomical organization (e.g. average connectivity and proportion of PYRs and INs), the simulated data was compared to PFC recordings.

To evaluate how accurately the different network architectures recapitulated the SUA statistics of the developing PFC, we employed the same parameters that we used to describe the experimental data. For each simulation we computed the simulated firing rates and STTC, and, for both variables, we extracted their skewness, kurtosis, Gini coefficient, plus the correlation between log-transformed firing rates and average STTC. We then derived the coordinates of the center of mass of the experimental data in this 7-dimensional parameter space, and calculated the its Euclidian distance from every simulation (see simplified scheme in Figure 5D).

Using this approach and multivariate linear regression, we found that the distance from the experimental data’s center of mass decreased as a function of the number of structural parameters that was set in its “extreme” configuration (log-normal distribution of synapse size and number, and correlated number of incoming and outgoing synapses) (Figure 5E). Further, the distance from the experimental center of mass decreased supra-linearly as a function of the number of “extreme” parameters (0-1 “extreme” parameters distance difference=0.07, 1-2 “extreme” parameters distance difference=0.09, 2-3 “extreme” parameters distance difference=0.12, linear model) (Figure 5E). This effect is also visually appreciable when the 7 dimensions in which we evaluated the model fit are reduced using tSNE (Supp. Figure 6A). When few synaptic parameters are set to their “extreme” configuration, there is little to no overlap between experimental and simulated data. Conversely, when all 3 synaptic parameters are set to their “extreme” configuration, the areas of maximum density of the two distributions are overlapping (Supp. Figure 6A).

**Figure 6.**
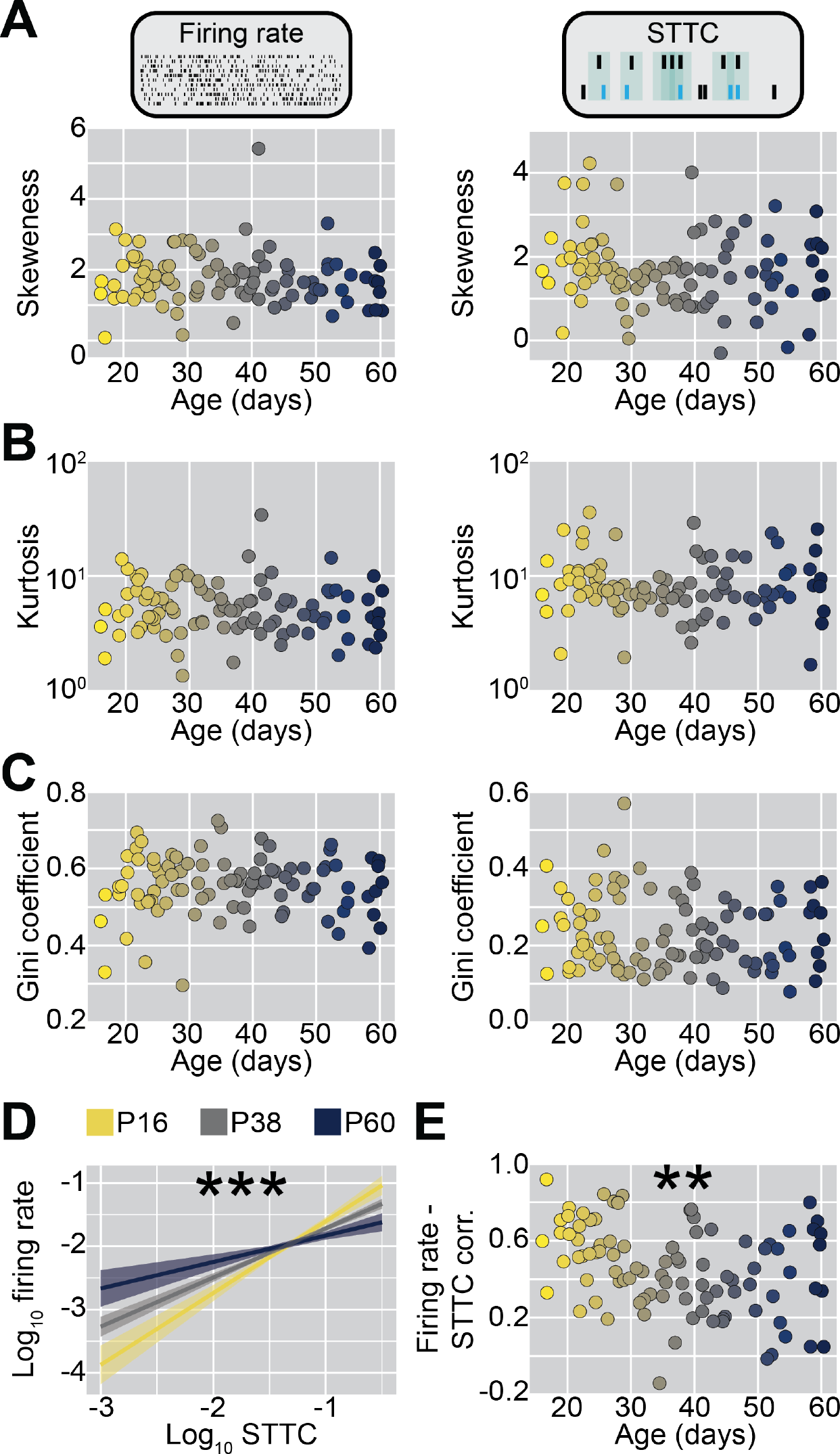
Skewness, kurtosis, Gini coefficient and correlations of firing rate and STTC in P16-60 mice. (**A**) Scatter plot displaying the skewness of firing rate (left) and STTC (right) of P16-60 mice (n=24 mice and 95 recordings). Color codes for age with 1-day increments. (**B-C**) Same as (**A**) for kurtosis and Gini coefficient. (**D**) Line plot displaying the log-transformed average STTC as a function of log-transformed firing rate across ages (n=24 mice and 2498 single units). Color codes for age. (**E**) Same as (**A**) for the Pearson correlation between log-transformed average STTC and log-transformed firing rate within individual mice. In (**A-C**) and (**E**) colored dots indicate individual recordings. In (**D)** data is presented as mean and 95% confidence interval. Asterisks in (**D-E**) indicate a significant effect of age. ***p<0.001, **p<0.01, linear mixed-effect models.

Taken individually, the structural parameter with the largest effect on the average distance from the experimental data’s center of mass was the distribution of synapse number (regression coefficient=-0.16, linear model), followed by the distribution of synaptic weight (regression coefficient=-0.06, linear model) and by the proportionality of incoming and outgoing synapses (regression coefficient=-0.06, linear model) (Figure 5F). Next, we considered the effect of the 3 synaptic parameters on individual properties of the simulated SUA statistics (Supp. Figure 6B-H). We found that the distribution of the synapse number had the largest effect on: (i-ii) skewness of firing rate and STTC, (iii) the kurtosis of STTC, (iv) the Gini coefficient of firing rate and (v) the log-log correlation between firing rate and average STTC. The proportionality of incoming and outgoing synapses had the largest effect on: (i) the kurtosis of firing rate and (ii) the Gini coefficient of STTC. The distribution of synaptic weights did not have the largest effect on any of the 7 parameters (Supp. Figure 6B-H).

**Supplementary Figure 6.**
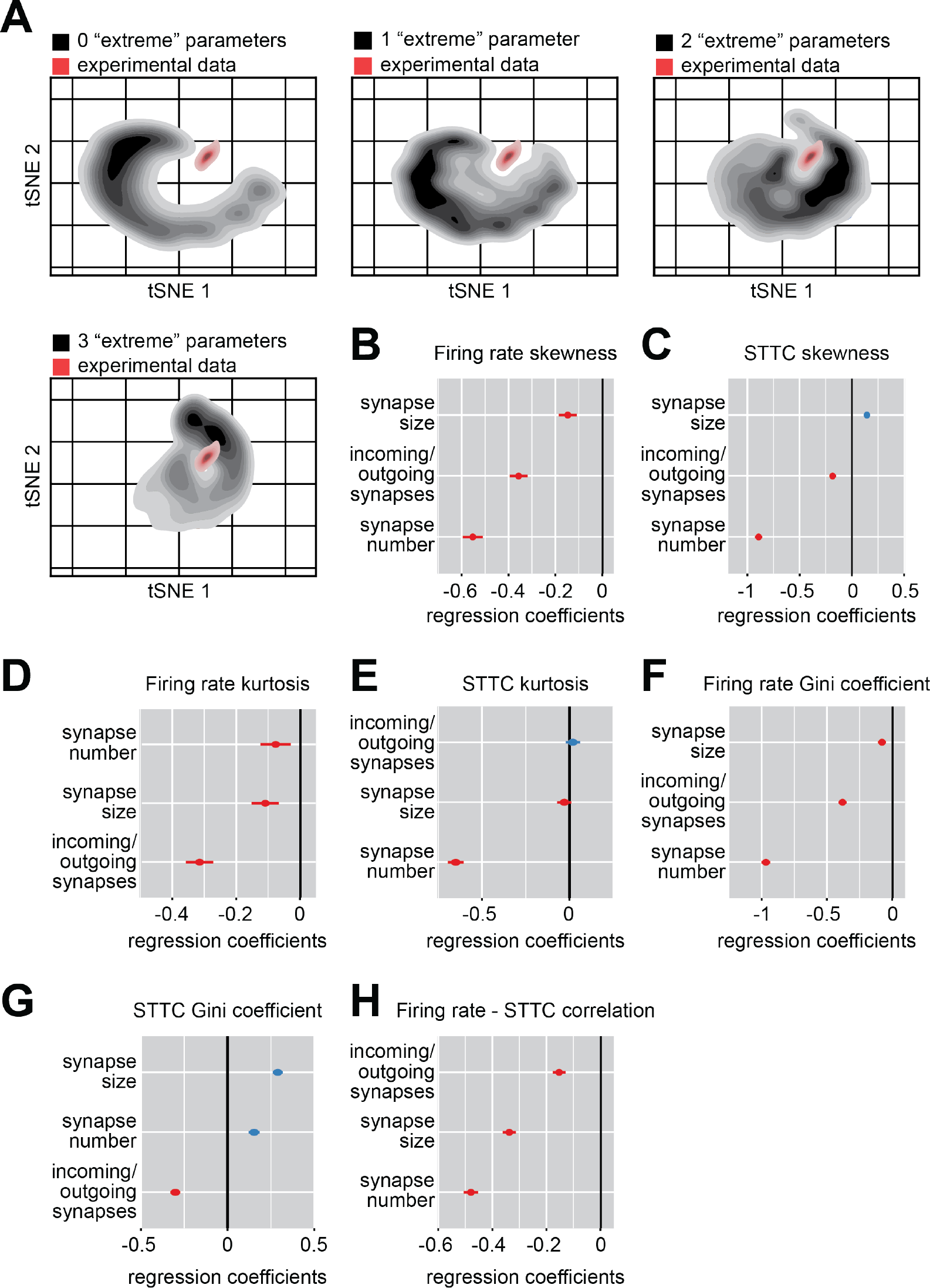
Spiking neural network modeling of the of the distribution shape of SUA statistics in the developing PFC. (**A**) tSNE-embedded space occupied by experimental (red) and simulated (black) data in the 7-dimensional parameter space used to evaluate model fit, as a function of the number of synaptic parameters set in their “extreme” configuration. Color intensity codes for probability. (**B**) Multivariate regression coefficients for the 3 synaptic parameters over the distance from the median firing rate skewness of experimental data. (**C-H**) Same as (**B**) for STTC skewness (**C**), firing rate kurtosis (**D**), STTC kurtosis (**E**), firing rate Gini coefficient (**F**), STTC Gini coefficient (**G**) and firing rate – STTC correlation (**H**). In (**B-H**) regression coefficients are presented as mean and 95% confidence interval.

Taken together, these data indicate that, in spiking neural networks simulations, extreme distributions of synaptic parameters are required to stably recapitulate the SUA statistics that we observed in the developing PFC. The distribution of synapse number on individual neurons is the synaptic parameter with the largest influence on the model fit. However, all three synaptic parameters play a significant role and influence different aspects of the simulated data.

### The shape of the SUA statistics distributions is stable throughout adulthood, while the oligarchy decreases with age

So far, we have shown that already shortly after birth the brain has a right-skewed, heavy-tailed and unequal distribution of SUA statistics that largely does not vary during the first two postnatal weeks. Since similar properties have been previously reported for the adult brain, the question arises if and how the extremeness of these parameters varies throughout late development and into adulthood. To address this question, we chronically recorded from the PFC of head-fixed mice from P16 to P60 (n=24 mice, 95 recordings, 2498 single units and 36899 spike train pairs).

SUA firing rate and STTC did not vary with age (age coefficient=-0.003 and - 10^-4^, CI [-0.003; 0.001] and [-6*10^-4^; 4*10^-4]^, p=0.28 and p=0.63, respectively, generalized linear mixed-effect model) (Supp. Figure 7A-B).

**Supplementary Figure 7.**
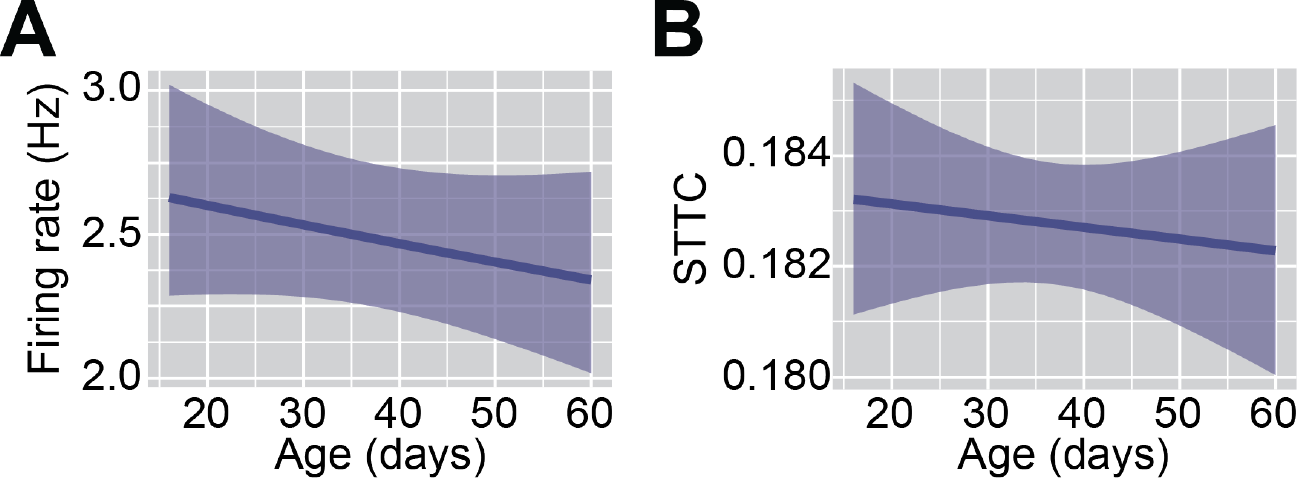
Prefrontal SUA firing statistics in P16-60 mice. (**A**) Line plot displaying the SUA firing rate of P16-60 mice (n=24 mice, 95 recordings and 2498 single units) as a function of age. (**B**) Same as (**A**) for STTC (n=24 mice, 95 recordings and 36899 spike train pairs). In (**A)** and (**B**) data is presented as mean and 95% confidence interval.

**Figure 7.**
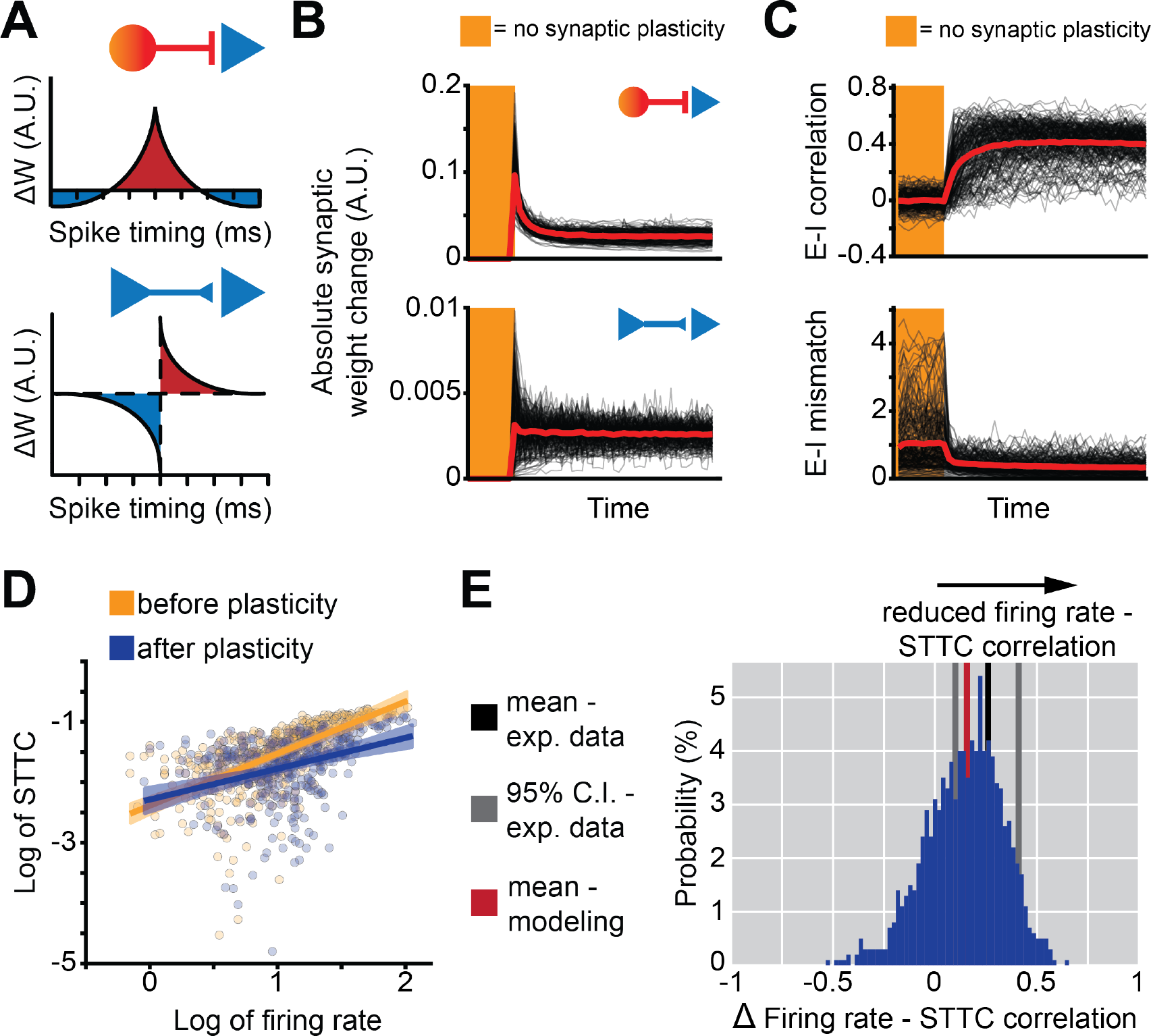
Spike-dependent inhibitory synaptic plasticity decreases the correlation between firing rate and STTC. (**A**) Schematic representation of the simulated inhibitory (top left) and excitatory (bottom left) synaptic plasticity rules. (**B**) Line plot of the IN-PYR (top) and PYR-PYR (bottom) synaptic weight changes over time. The shaded area indicates the period without synaptic plasticity. (**C**) Line plot of excitatory and inhibitory currents correlation (top) and absolute difference (bottom) over time. The shaded area indicates the period without synaptic plasticity. (**D**) Scatter and line plot of a representative example of the correlation between simulated firing rate and average STTC before (yellow) and after (blue) synaptic plasticity. (**E**) Histogram plot of the difference in simulated firing rate and average STTC correlation before and after synaptic plasticity. The black line indicates the mean of the experimental data, the two grey lines the 95% C.I. of the mean and the red line the mean of the simulated data. In (**B-C**) individual black lines represent individual simulations and the red line the mean across simulations. For visualization purposes, only 1/5 of the individual simulations are shown. In (**D**) individual dots represent individual simulated neurons, and data is presented as mean and 95% C.I.

Analogously to the data for the neonatal brain, the skewness of the SUA statistics did not change also for the P16-60 developmental time window (age coefficient=-0.004 and −0.01, CI [-0.02; 0.007] and [-0.02; 0.003], p=0.49 and p=0.14, for firing rate and STTC, respectively, linear mixed-effect model) (Figure 6A). Along the same line, also the firing rate’s and STTC’s kurtosis (age coefficient=-0.001 and - 0.001, CI [-0.005; 0.002] and [-0.005; 0.002], p=0.42 and p=0.55, for firing rate and STTC, respectively, linear mixed-effect model) and Gini coefficient (age coefficient=10^-5^ and 7*10^-4^, CI [-0.001; 0.001] and [-0.002; 8*10^-4^], p=0.99 and p=0.34, for firing rate and STTC, respectively, linear mixed-effect model) remained constant for the investigated time window (Figure 6B-C).

Similar to the data for early development, in the adult PFC, the log-transformed firing rate of a neuron and its average STTC correlated with each other (average STTC coefficient=1.53, CI [1.34; 1.71], p<10^-56^, linear mixed-effect model) (Figure 6D). However, when we fitted a model that includes an interaction of STTC with age, the correlation strength between the two variables decreased with age (age and average STTC interaction coefficient=-0.03, CI [-0.05; −0.02], p<10^-5^, linear mixed-effect model) (Figure 6D). Accordingly, the Pearson correlation between firing rate and average STTC on an individual mouse basis strongly decreased with age, from roughly ∼0.6 to ∼0.3 (age coefficient=-0.006, CI [-0.009; −0.002], p=0.001, linear mixed-effect model) (Figure 6E).

Thus, in the mouse PFC, the extremeness of SUA statistics distributions does not significantly change between P16 and P60. However, the correlation between firing rate and STTC is roughly halved from P16 to P60, indicating that the oligarchical state in PFC is more pronounced during early development than at adulthood.

### Inhibitory synaptic plasticity parsimoniously explains the gradual disappearance of the oligarchy

Throughout the same developmental phase during which we observed a decrease of the correlation between the firing rate of a neuron and its average STTC, the cortex transitions into a state of detailed E-I balance^55^, a phenomenon that has been linked to inhibitory synaptic plasticity^56^. Given the critical role that E-I ratio plays in controlling correlations among neurons^38^, we hypothesized that this transition might explain the progressive decline of the correlation between firing rate and STTC.

To test this hypothesis, we resorted to spiking neural network modeling with an analogous architecture as the one used to model early cortical activity. After an initial period in which the networks were run with frozen synaptic sizes and all the synaptic parameters in their “extreme” configuration, we added symmetric spike-time dependent inhibitory plasticity (ISTDP) on the synapses connecting INs to PYRs^56^ (Eq. 1) and classic asymmetric Hebbian plasticity (STDP) on PYR-PYR excitatory synapses^57^ (Figure 7A). The original formulation of the ISTDP rule implemented a single target firing rate (“α”, equation 1) for the entire population of PYRs. In this study, we instead decided to draw “α” from a lognormal distribution, to better recapitulate the diversity of the experimentally observed firing rates (see Materials and Methods for details):

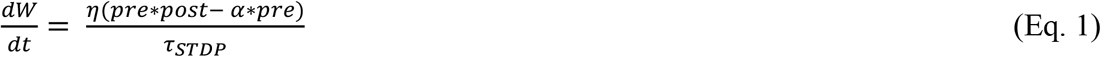

where *W* is an IN-PYR synaptic weight, *pre* and *post* are the pre- and postsynaptic activity, *α* is the target rate for the postsynaptic PYR, drawn from a log-normal distribution, *τ*_*STDP*_ is the decay time constant of the plasticity rule, and *η* is the learning rate.

Analogously to the approach that we previously described, also for this set of simulations, we treated several other structural parameters as random variables, and we systematically varied them within the same biologically-constrained range of values that we employed for the former simulations. To evaluate the effect of synaptic plasticity, we compared the SUA statistics of the frozen synapses phase and those of the last fifth of the simulated data, a phase in which synaptic changes were stable (Figure 7B).

Analogously to previous results^56^, we found that adding ISTDP resulted in an increase of correlation between incoming excitatory and inhibitory currents across neurons (Figure 7C, top) and a decrease of the mismatch (i.e. the absolute value of the difference) between excitatory and inhibitory currents across time (computed in 5s bins) within individual neurons (Figure 7C, bottom).

In line with our hypothesis, we found that adding ISTDP and STDP to the network also induced a decrease in the correlation between firing rate and average STTC that was comparable to what we experimentally observed in P16-60 mice (mean decrease in firing rate – average STTC correlation in simulated data=0.16, in experimental data=0.26, CI [0.11; 0.41], Figure 7D-E). When we further evaluated the effects of synaptic plasticity, we found that it only minimally affected the skewness, kurtosis and Gini coefficient of firing rate and STTC, similar to the experimental data from the PFC of P16-60 mice (Supp. Figure 8). Of these 6 parameters, only the kurtosis and Gini coefficient of firing rate were narrowly outside the 95% confidence interval of the experimental values (Supp. Figure 8).

**Supplementary Figure 8.**
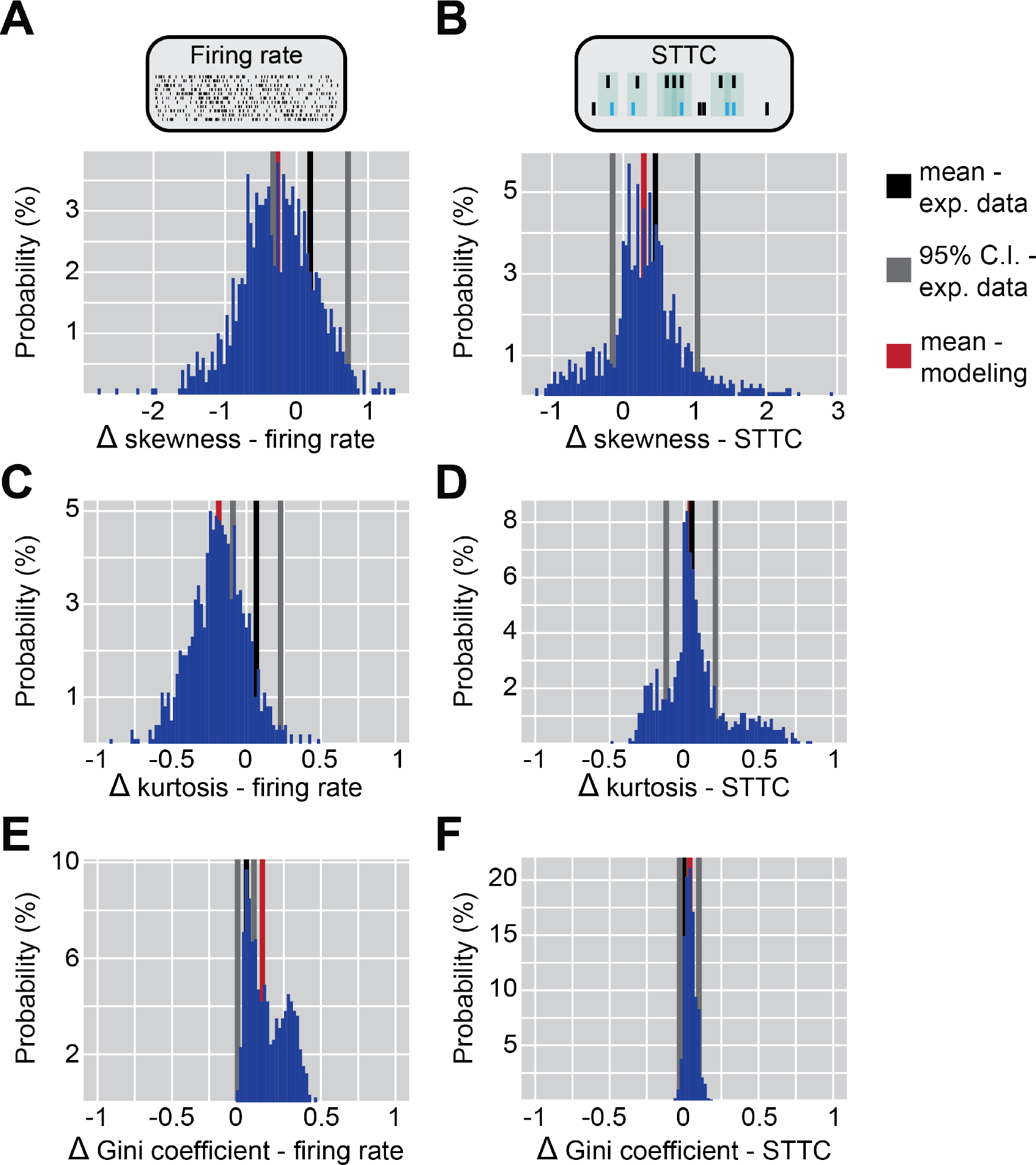
Synaptic and network effects of synaptic plasticity rules. (**A**) Histogram plot of the difference in simulated skewness of firing rate before and after synaptic plasticity. The black line indicates the mean of the experimental data, the two grey lines the 95% C.I. of the mean and the red line the mean of the simulated data. (**B-F**) Same as (**A**) for the skewness of STTC (**B**), kurtosis of firing rate (**C**) and STTC (**D**) and Gini coefficient of firing rate (**E**) and STTC **(F**).

Taken together, these data indicate that adding ISTDP and STDP to a spiking neural network parsimoniously recapitulates the developmental decrease in the correlation between firing rate and STTC while only minimally affecting the distributions of firing rate and STTC, consistent with what we observed in the PFC of P16-60 mice.

## Discussion

A fascinating property of the adult brain is that a large number of parameters, ranging from the size of synapses to the power of extracellular currents^1,3,4^, follow extreme distributions. This organization has been suggested to have many desirable properties^10,27^, but how and when it arises is still a matter of debate. Here, we report that in the PFC and OB, two brain regions exemplifying slow and fast maturational dynamics, already in the first days of extrauterine life, the distributions of first- and second-order SUA statistics are extreme: right-skewed, heavy-tailed and highly unequal. While the central tendency of these distributions varies across development, their shape remains largely unchanged until adulthood. We also show that early brain activity is in an oligarchical state, in which high firing rate neurons display hub-like properties and exert a disproportionate influence on their local network, a phenomenon that becomes less prominent as mice age. Leveraging spiking neural network modeling we demonstrate that, to recapitulate these network properties, analogously extremely distributed synaptic parameters are needed. We conclude by showing that the progressive disappearance of the oligarchical state can be parsimoniously explained by introducing an inhibitory synaptic plasticity rule that establishes a detailed excitation-inhibition balance. This work suggests that the distribution shape of structural and functional neural parameters is not fundamentally altered by developmental processes, but rather preconfigured and experience-independent.

In altricial animals such as rodents, the first postnatal week roughly corresponds to mid-late gestation in humans^37^. At this early stage, brain activity has several unique traits, such as discontinuity^37^, highly correlated spiking activity^38,42,43^, the presence of transient cell types and circuits^58^, extremely low firing rates^38,59^, low levels of inhibition^38,60,61^ and a loose temporal coordination of excitation and inhibition^55,56^. Importantly, sensory systems are still very underdeveloped. In rodents, the retina becomes light sensitive around P8-9^59,62^, and eye opening only takes place around P14-P15, a few days after hearing onset occurs^63^. The whisking-related sensory system also follows a similar timeline. In the first postnatal week, ∼90% of whisker movements do not induce increased firing rate in the somatosensory cortex, and ∼90% of firing in the somatosensory cortex is unrelated to whisker movements^64^. Even after splitting whisker movements by size and only considering the few large ones that occur, the somatosensory cortex is active concomitantly with a whisker movement less than 50% of the time^65^. Whisker-elicited sensory responses are initially mainly the result of passive stimulations by the dam and the littermates, while robust active whisking only emerges around P10-12^66–68^. Solely the olfactory system follows a distinct developmental dynamic and, even though it is also still developing^69^, it is already functional and behaviorally relevant in the first postnatal days^35,36^. Despite the paucity of sensory information that is therefore available in the first postnatal week, we find that the distribution of SUA statistics is already extreme and does not significantly vary from early development to adulthood. Corroborating the idea that experience does not play a significant role in this process, we find no differences between the OB, that is already “online”, and the PFC, which is supposed to be one of the slowest developing brain regions^37^. We further show that already from P4 onwards, the PFC also displays a complex network topology that is reminiscent of small-world networks, a property that is typical of many real world networks^70^, including the adult brain^48–50^ (but see^51^).

These results are difficult to reconcile with the hypothesis that brain structure is initially “diffuse” and only later acquires the structural organization that is typical of the adult brain^29,30,71^. Rather, they support several recent studies that have highlighted the importance of “nature over nurture”^72,73^ in establishing neural circuits and the distributions of their parameters. For instance, in neuronal cultures, the variance of glutamatergic synaptic sizes does not differ between networks that are silenced from plating onwards, and networks that are spontaneously active^7^. Similarly, cultures of dissociated neurons exhibit the ability to self-organize into a complex network topology with small-world properties that is characterized by extreme distributions of firing rates and connection weights^47^. Extreme distributions of firing rates and functional connectivity are also observed in human brain organoids^74,75^, that also do not have access to sensory inputs. Along the same lines, *in vivo* blocking of synaptic transmission in the hippocampus of anesthetized rats does not alter the distribution of spine sizes^76^. Even more surprisingly, completely abolishing all central nervous system activity for the first four days of life of the larval zebrafish only minimally impacts the tuning and functionality of neurons, and the ability of the fish to learn a complex visuomotor task^72^. Intriguingly, similar concepts are beginning to percolate also in the field of artificial intelligence^77^, which has been traditionally dominated by a bottom-up and learning-based approach.

The relationship between the firing rate of a neuron and the strength of its pairwise interactions with other neurons is complex and still debated. A number of experimental and theoretical studies found a positive correlation between the two variables^78–81^. However, this relationship has also been reported to be absent or even negative^82–85^. Here we show that, in the PFC and OB of P4-12 mice, there is a strong positive correlation between firing rate and average pairwise interaction of a neuron, as measured by the STTC coefficient. Thus, neurons with extreme firing rates are also likely to have extreme average STTC values, an organization that we refer to as oligarchical. The correlation between firing rate and STTC weakens throughout development, but it is present also in adulthood. In the early PFC, firing rate also correlates with a neuron’s “hubness score”, a composite metric that encompasses 5 different measures of local and global hubness. The topic of hub neurons, defined as a subclass of neurons that has an outsized influence on the network activity, has been the subject of extensive experimental and theoretical research in the developing brain^52,86– 88^. In agreement with our results, hub neurons in the developing entorhinal^86,87^ and barrel cortex^52^ are also characterized by high firing rates and high functional connectivity, a prediction that is shared by theoretical work^89^. Previous work has also shown that hub neurons are generally INs, something that we could not investigate in the current study, due to the difficulty of separating INs and PYRs based on their waveform properties in the early developing brain^90^. This work further shows that, in a neural network model, adding inhibitory synaptic plasticity results in E-I balance across neurons and time, and decreases the influence of high firing rate neurons on the network activity. Thus, we propose that the loose temporal E-I balance that is typical of the developing brain^55,56^ might be a permissive mechanism for the role exerted by hub neurons on their surroundings. Whether the developmental shift of E-I ratio towards inhibition^38,60^ also plays a role remains to be investigated.

Lastly, we show that there is a mechanistic link between the distribution of structural (synaptic) and functional (SUA statistics) parameters. In particular, we report that, in a spiking neural network model, three synaptic parameters had a strong influence on the extremeness of the simulated spiking activity: (i) whether the size of synapses followed a log-normal distribution, (ii) whether the number of synapses of individual neurons followed a log-normal distribution, and (iii) whether the number of dendritic and axonic (incoming and outgoing) synapses were correlated with each other. The three parameters had a synergistic effect on the model goodness of fit to the experimental data, and had a distinct impact on different properties (skewness, kurtosis and Gini coefficient) of the simulated firing rate and STTC distributions. While a large body of evidence supports the notion that the size and number of synapses follow an extreme distribution^1^, to the best of our knowledge, whether the number of dendritic and axonic synapses are correlated with each other has not been explicitly investigated in mammals. However, a recent study found that, in the mouse PFC, the length of a neuron’s dendrite and axon are positively correlated^91^. Further, in a full-brain Drosophila Melanogaster reconstruction, the number of incoming and outgoing synapses are tightly correlated with each other (Person coefficient = 0.8)^92^. The fact that extremely distributed synaptic parameters are required to faithfully reproduce the SUA statistics of the developing PFC suggests that the distribution of synaptic parameters is experience-independent.

This study has several limitations. First, despite investigating mice from a very early developmental phase, and doing so also in the PFC, one of the slowest developing brain regions, we cannot exclude that some experience-dependent processes have not already taken place. Second, we do not directly experimentally probe whether the distribution shape of synaptic parameters is stable across the first postnatal weeks. However, our modeling results generate a number of predictions that could be experimentally addressed: (i) that synapse size and number on individual neurons follow extreme distributions already in the first postnatal week, (ii) that the shape of these distributions should be stable across development, and (iii) that the number of incoming and outgoing synapses should be tightly correlated. Third, while our results argue against a role of experience in shaping these processes, we do provide an alternative mechanistic answer to the question. This topic has however already been the subject of a number of theoretical studies^7,9,76,89^. Further, we generated a large and detailed experimental open-access database that could be instrumental in benchmarking future research on this topic. An approach that we believe might be of particular interest is that of generative models^93,94^. Illustrating the promise of this approach, generative models solely based on spatiotemporal gradients of neuronal development^95^ or homophily principles^75^ can recapitulate important features of the complex topology of adult brains.

In summary, we report that the extremeness with which functional and structural parameters are distributed is stable across a large portion of the lifespan, which suggests that the brain is in a preconfigured state, and that experience-dependent processes do not fundamentally alter its organization. Further elucidating the principles underlying the establishment of neural circuits might prove insightful for advancing the field of biological and artificial intelligence alike.

## Acknowledgments

We thank Sebastian Bitzenhofer and Irina Pochinok for valuable discussions and feedback on the manuscript, P. Putthoff, A. Marquardt and A. Dahlmann for excellent technical assistance. This work was funded by grants from the European Research Council (ERC-2015-CoG 681577 to I.L.H.-O.), Marie Curie Training Network euSNN (MSCA-ITN-H2020-860563 to I.L.H.-O.), Horizon2020 DEEPER 101016787, the German Research Foundation (437610067, 178316478 and 302153259 to I.L.H.-O.) and Landesforschungsförderung Hamburg (LFF76, LFF73 to I.L.H.-O.).

## Author Contributions

M.C. and I.L.H.-O. designed the experiments and wrote the manuscript. M.C., J.K.K., M.H. and Y.-N.C. carried out the experiments, M.C. analyzed the experimental data and carried out neural network modeling. All authors interpreted the data, discussed and commented on the manuscript.

## Data and code availability

SUA data that was newly generated for this study will be available at the following open-access repository: https://gin.g-node.org/mchini/Chini_et_al_Preconfigured Code, processed data and statistical analysis supporting the findings of this study will be available at the following open-access repository: https://github.com/mchini/Chini_et_al_Preconfigured

### Experimental models and subject details

All experiments were performed in compliance with the German laws and following the European Community guidelines regarding the research animal’s use. All experiments were approved by the local ethical committee (G132/12, G17/015, N18/015, N19/121). Experiments were carried out on C57BL/6J mice of both sexes. Mice were housed in individual cages on a 12 h light/12 h dark cycle, and were given access to water and food ad libitum. P16-60 mice were housed with a minimum of two cage-mates after weaning. The day of birth was considered P0. Details on the data acquisition and experimental setup of open-access datasets that were used in this project have been previously published^38,39^.

### *In vivo* electrophysiology in P4-12 mice

#### Surgery

In vivo extracellular recordings were performed from the PFC and the ventral portion of the OB of non-anesthetized P4-P12 mice. The surgery and animal preparation were analogous for the two brain regions. Before starting with the surgical procedure, a local anesthetic was applied on the neck muscles (0.5% bupivacain / 1% lidocaine). The surgery was performed under isoflurane anesthesia (induction: 5%; maintenance: 1-3%, lower for older pups, higher for younger pups). Neck muscles were severed to minimize muscle artifacts. A craniotomy over the PFC (0.5 mm anterior to bregma, 0.1-0.5 mm lateral to the midline) or the OB (0.5–0.8 mm anterior to frontonasal suture, 0.5 mm lateral to inter-nasal suture) was performed by carefully thinning the skull and then removing it with the use of a motorized drill. Mice were head-fixed into a stereotactic frame and kept on a heated (37°) surface surrounded by cotton wool throughout the entire recording. To record from the PFC, a Neuropixels probe 1.0 phase 3B (Imec, Belgium) was slowly vertically inserted (angle 0°) into the frontal lobe (insertion time 20-30 minutes), at a depth varying between 2.6 and 4 mm depending on the age of the animal. Due to the small size of the brain, in younger animals, not all 384 recording channels were inserted in the brain. The tip of the probe was used as reference. To record from the OB, a single-shank silicon probe (NeuroNexus, MI, USA) with 16 recording sites and 50 μm inter-site spacing was vertically inserted (angle 0°) at a depth varying between 1.0 and 2.0 mm. A silver wire inserted in the cerebellum was used as reference. Both probe types were inserted using a micromanipulator. Before signal acquisition, mice were allowed to recover for ∼45 minutes, to maximize the quality and stability of the recording as well as single units’ yield.

#### Signal acquisition

For PFC recordings, signals from the bottom 384 channels were recorded at a 30 kHz using the Neuropixels head-stage 1.0 and Neuropixels 1.0 PXIe acquisition system (Imec, Belgium). The SUA signal was acquired through the OpenEphys interface and the Neuropixels plugin (AP gain = 500, AP Filter Cut = ON).

For OB recordings, signals were band pass-filtered (0.1–9 kHz) and digitized (32 kHz) by a multichannel amplifier (Digital Lynx SX; Neuralynx, Bozeman, MO, USA) and acquired through the Cheetah acquisition software (Neuralynx, Bozeman, MO, USA).

#### Histology

Epifluorescence images of coronal brain sections were acquired post mortem to reconstruct the position of the DiI-stained recording electrode. Only mice in which the electrodes were placed in the correct position were kept for further analysis.

### *In vivo* electrophysiology in P16-60 mice

#### Surgery

Acute In vivo extracellular recordings were performed from the PFC of awake, head-fixed mice of both sexes. Before starting with the surgical procedure to implant a metal head-plate (Neurotar, Helsinki, Finland) for head-fixation, buprenorphine (0.5 mg/kg bw) was injected subcutaneously. The surgery was performed under isoflurane anesthesia (induction: 5%; maintenance: 2.5%). Anesthesia depth was confirmed with the paw withdrawal reflex. Eyes were covered with an ointment (Vidisic, Bausch + Lomb, Berlin, Germany) to prevent them from drying out. After disinfection with Betasisodona, the scalp was removed from the top of the head and the edges treated for analgesia with application of a Lidocain/Bupivicain mixture (0.5% bupivacain / 1% lidocaine). A craniotomy was performed to make the mPFC (0.5–2.0 mm anterior to bregma, 0.1–0.5 mm right to the midline) accessible for recordings. A synthetic window was fixed to the skull around the craniotomy to be able to protect the tissue with Kwik-Cast sealant (World Precision Instruments, Friedberg, Germany). A silver wire, serving as a ground and reference electrode, was inserted between the skull and cerebellum. The metal head-plate was attached to the skull with dental cement. For recovery from anesthesia, mice were placed in a cage on a heating mat and after being fully awake, they were put back into their home cage with their cage mates. For further analgesia Metacam (0.5 mg/ml, Boehringer-Ingelheim, Germany) was mixed into soft food and provided for 48 h after the surgery.

#### Training and signal acquisition

After recovery from the surgery, mice were accustomed to the head-fixation system and trained to move the air-lifted carbon cage from the MobileHomeCage system (Neurotar, Helsinki, Finland). To perform electrophysiological recordings, the craniotomy was uncovered and an electrode (NeuroNexus, MI, USA) was stereotactically inserted into the mPFC (one-shank, A1×16-channel, 100 μm-spaced, 2.0 mm deep). The signal was acquired for 30-40 min. Extracellular signals were band-pass filtered (0.1–9000 Hz) and digitized (32 kHz) with a multichannel extracellular amplifier (Digital Lynx SX; Neuralynx, Bozeman, MO, USA).

#### Histology

Epifluorescence images of coronal brain sections were acquired post mortem to reconstruct the position of the recording electrode. To this aim, after the last recording, a DiI-coated electrode was inserted. Only mice in which the electrodes were placed in the correct position were kept for further analysis.

### Spike sorting

PFC recordings were spike-sorted with Kilosort 2.5^96^ (fshigh = 500, minFR = 0.001, spkTh = −4, sig = 20, nblocks = 5). OB recordings were spike-sorted with Klusta^97^.

Automatically-obtained clusters were manually curated using phy (https://github.com/cortex-lab/phy).

### SUA firing statistics and shape distribution parameters

#### Firing rate

Firing rate (in Hz) was computed as the number of spikes divided by the total recording length in seconds.

#### Spike-Time Tiling Coefficient (STTC)

The STTC (timescale of 10 ms) was computed as previously described^38,40^.

Briefly:

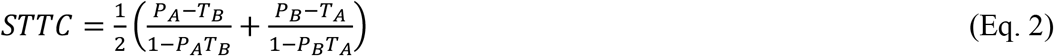

where P_A_ is the proportion of spikes of spike train A that occurs within ±Δt of a spike train B spike. T_A_ is the proportion of time that occurs within (is “tiled” by) ±Δt from spikes of spike train A. The same applies for P_B_ and T_B_. ±Δt is the lag parameter and was set at 10 ms.

#### Skewness

Skewness was computed using the homonymous Matlab function *skewness* as:

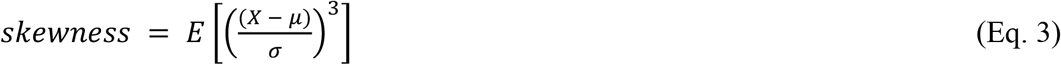

Where X is the random variable of interest, μ is its mean, σ its standard deviation, and E is the expectation operator.

#### Kurtosis

Kurtosis was computed using the homonymous Matlab function *kurtosis* as:

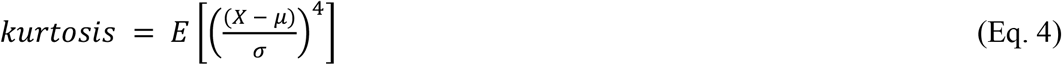

Where X is the random variable of interest, μ is its mean, σ its standard deviation, and E is the expectation operator.

#### Gini coefficient

The Gini coefficient was computed using the Matlab function *gini*^98^. The Gini coefficient is calculated by taking the ratio of the area that lies between the line of equality and the Lorenz curve, over the total area under the line of equality.

### Complex network properties

To calculate the network properties of the developing brain, we utilized symmetrical STTC matrices. The pre-processing consisted of thresholding and binarization. To threshold the data, we computed surrogate spiking data for each mouse individually. To account for the slow co-modulation of firing rates that is typical of the developing brain, we generated surrogate spiking data by leaving the timing of spikes unaffected and shuffling the identity of the neuron that emitted the spike. This pseudo-randomization thus generated surrogate spike vectors while preserving the population rate. Using these spike vectors, we then computed at least 1000 STTC values on shuffled data, and used the 90^th^ percentile value to threshold and binarize the real-data STTC matrices. The binary and undirected matrices were then analyzed with the MATLAB Brain Connectivity Toolbox^46^.

The clustering coefficient and transitivity of the matrices were extracted with the *clustering_coef_bu* and *transitivity_bu* functions, respectively. To compute the characteristic path length, we first computed the distance matrix (*distance_bin*) and then computed the characteristic path length (*charpath*) without including distances on the main diagonal and infinite distances. All these values were normalized by dividing them with a corresponding “null” value that was computed by generating 100 synthetic random networks (*makerandCIJ_und*), extracting the same parameters for each iteration, and computing the average. Small-worldness was computed by dividing the normalized clustering coefficient by the normalized characteristic path length.

The four “hubness” parameters that we used to compute the composite “hubness score” were also extracted with the same MATLAB toolbox. For each node individually, we computed its total amount of edges (i.e. its degree, using *degrees_und*), its total amount of weights (*strengths_und*, this is the only parameter that was computed on weighted and not binary matrices), its betweenness centrality (*betweenness_bin*, after using *weight_conversion* and ‘lengths’) and its closeness centrality (the inverse of the node’s average distance computed with *distance_bin*).

### Neural network modeling

All simulations were performed using Brain2 for Python3.7^99^.

#### General neural network architecture

The architecture of the network was set similarly to previously published work^38,100^, and is schematically illustrated in Figure 6A-C . The network was composed of 400 conductance-based leaky integrate-and-fire neurons, 80% of which were excitatory (PYRs) (N=320) and 20% were inhibitory (INs) (N=80). PYRs were simulated with outgoing excitatory synapses and INs with outgoing inhibitory synapses. Excitatory (PYR→PYR, PYR→IN) and inhibitory (IN→IN and IN→PYR) synapses were simulated as AMPA and GABA conductances, respectively. Due to the near-instantaneous rise times of AMPA- and GABA-mediated currents (both typically <0.5 ms), we opted to neglect these in the simulations. Synaptic transmission was assumed to be instantaneous (i.e. with zero delay).

The dynamics of each excitatory and inhibitory cell were governed by the following stochastic differential equation:

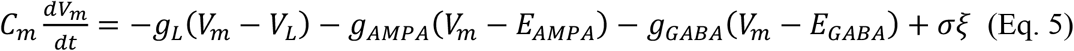

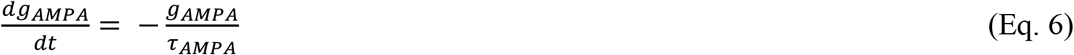

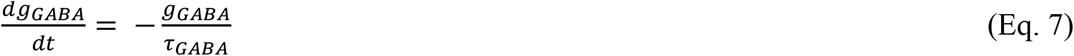

where *V*_*m*_ is the membrane potential, *V*_*L*_ is the leak membrane potential and *E*_*AMPA*_ and *E*_*GABA*_denote the AMPA and GABA current reversal potentials, respectively. The synaptic conductance parameters and the corresponding decay time constants are denoted by *g*_*AMPA*_, *g*_*GABA*_and *τ*_*AMPA*_, *τ*_*GABA*_, respectively. *σ* ξ is a noise term that is generated by an Ornstein-Uhlenbeck process with zero mean. The networks were simulated for a duration of 10 s (simulations without plasticity) or 50 s (simulations with plasticity). All simulations were performed with a time step (dt) of 0.1 ms and integrated with Euler’s method. All parameter values/ranges used in the simulations are listed in Table 1.

**Table 1.**
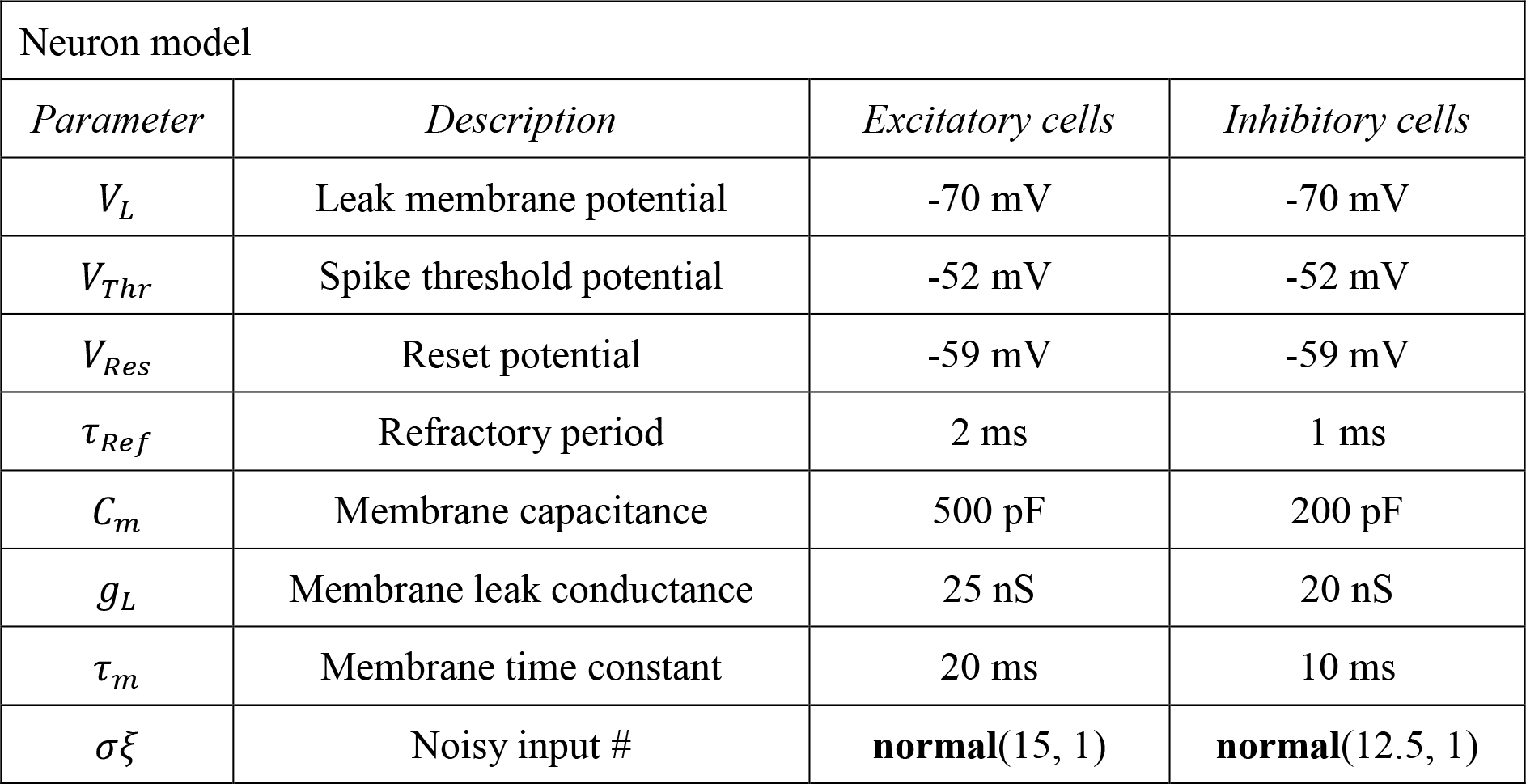
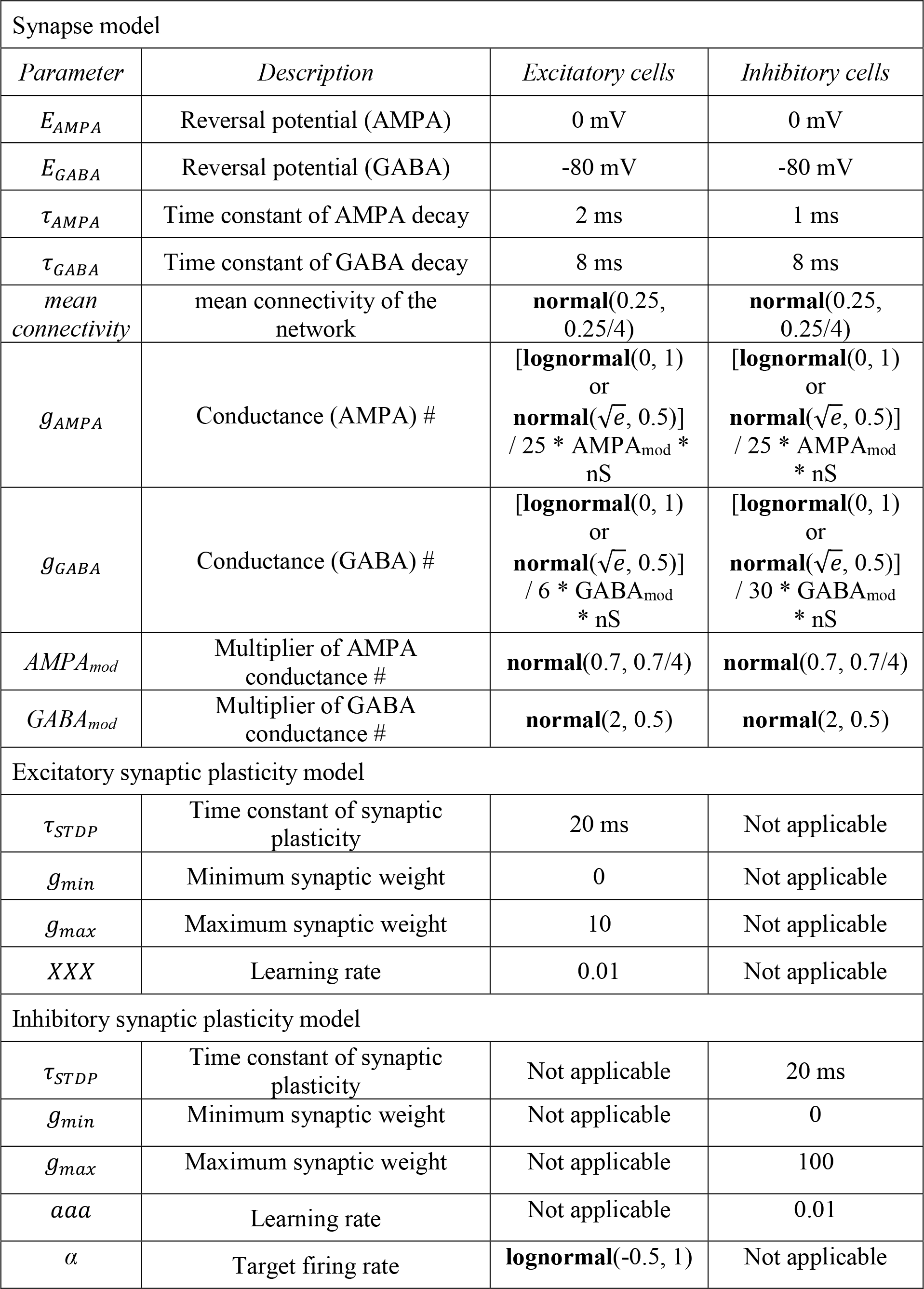
Parameters of the leaky integrate-and-fire network. “Normal” and “lognormal” refer to values of variables that are randomly drawn from a normal or lognormal distribution, respectively. The two values in parenthesis refer to, respectively, the mean and the standard deviation of the (underlying) normal distribution. ^#^Note that the two distributions have the same mean (i.e. the mean of lognormal(0, 1) is 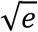).

| Neuron model |  |  |  |
| --- | --- | --- | --- |
| *Parameter* | *Description* | *Excitatory cells* | *Inhibitory cells* |
| $V_L$ | Leak membrane potential | -70 mV | -70 mV |
| $V_{Thr}$ | Spike threshold potential | -52 mV | -52 mV |
| $V_{Res}$ | Reset potential | -59 mV | -59 mV |
| $\tau_{Ref}$ | Refractory period | 2 ms | 1 ms |
| $C_m$ | Membrane capacitance | 500 pF | 200 pF |
| $g_L$ | Membrane leak conductance | 25 nS | 20 nS |
| $\tau_m$ | Membrane time constant | 20 ms | 10 ms |
| $\sigma\xi$ | Noisy input # | **normal(15, 1)** | **normal(12.5, 1)** |

**Table 1.** Parameters of the leaky integrate-and-fire network. “Normal” and “lognormal” refer to values of variables that are randomly drawn from a normal or lognormal distribution, respectively. The two values in parenthesis refer to,
| Synapse model |  |  |  |
| --- | --- | --- | --- |
| *Parameter* | *Description* | *Excitatory cells* | *Inhibitory cells* |
| $E_{AMPA}$ | Reversal potential (AMPA) | 0 mV | 0 mV |
| $E_{GABA}$ | Reversal potential (GABA) | -80 mV | -80 mV |
| $\tau_{AMPA}$ | Time constant of AMPA decay | 2 ms | 1 ms |
| $\tau_{GABA}$ | Time constant of GABA decay | 8 ms | 8 ms |
| *mean connectivity* | mean connectivity of the network | **normal**(0.25, 0.25/4) | **normal**(0.25, 0.25/4) |
| $g_{AMPA}$ | Conductance (AMPA) # | [**lognormal**(0, 1) or **normal**($\sqrt{e}$, 0.5)] / 25 * AMPA<sub>mod</sub> * nS | [**lognormal**(0, 1) or **normal**($\sqrt{e}$, 0.5)] / 25 * AMPA<sub>mod</sub> * nS |
| $g_{GABA}$ | Conductance (GABA) # | [**lognormal**(0, 1) or **normal**($\sqrt{e}$, 0.5)] / 6 * GABA<sub>mod</sub> * nS | [**lognormal**(0, 1) or **normal**($\sqrt{e}$, 0.5)] / 30 * GABA<sub>mod</sub> * nS |
| $AMPA_{mod}$ | Multiplier of AMPA conductance # | **normal**(0.7, 0.7/4) | **normal**(0.7, 0.7/4) |
| $GABA_{mod}$ | Multiplier of GABA conductance # | **normal**(2, 0.5) | **normal**(2, 0.5) |
| Excitatory synaptic plasticity model |  |  |  |
| $\tau_{STDP}$ | Time constant of synaptic plasticity | 20 ms | Not applicable |
| $g_{min}$ | Minimum synaptic weight | 0 | Not applicable |
| $g_{max}$ | Maximum synaptic weight | 10 | Not applicable |
| XXX | Learning rate | 0.01 | Not applicable |
| Inhibitory synaptic plasticity model |  |  |  |
| $\tau_{STDP}$ | Time constant of synaptic plasticity | Not applicable | 20 ms |
| $g_{min}$ | Minimum synaptic weight | Not applicable | 0 |
| $g_{max}$ | Maximum synaptic weight | Not applicable | 100 |
| aaa | Learning rate | Not applicable | 0.01 |
| $\alpha$ | Target firing rate | **lognormal**(-0.5, 1) | Not applicable |

#### Random variables

To avoid choosing an arbitrary neural network architecture, a number of parameters were treated as “random variables” and systematically varied across a biologically-constrained range. There parameters included: 1) the noisy input (Ornstein-Uhlenbeck process) that was used to independently drive PYRs and INs; 2) the mean probability with which neurons were connected, which was the same for all possible population combinations: i) PYR-PYR, ii) PYR-IN, iii) IN-PYR, iv) IN-IN; 3) the mean size of excitatory and inhibitory synaptic weights.

#### Synaptic parameters

In these networks, we studied the influence on the simulated SUA statistics exerted by three synaptic parameters: whether the size of synapses followed a normal or a log-normal distribution, whether the number of synapses of individual neurons followed a normal or a log-normal distribution, and whether the number of dendritic and axonic (incoming and outgoing) synapses were correlated or uncorrelated with each other. The 2 possible configurations for the 3 synaptic parameters resulted in 2^3^=8 different network types, of which we simulated 1000 each. These 3 synaptic parameters were simultaneously varied for the entire network and not in a neuronal population-specific manner.

#### Number of synapses and correlation between dendritic and axonic synapses

We first generated a normal (mean = 1, std = 1/4) or log-normal (mean of the underlying normal

= 0, std of the underlying normal = 0.5) relative distribution of the number of connections that was normalized to have a sum=1. To account for the different number of connections in different simulations (see previous paragraph *Random variables*), the relative distribution was then scaled by the total amount of connections between the pre- and post-synaptic population according to:

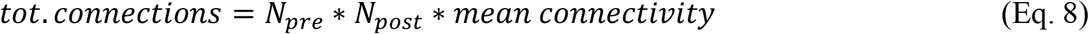

Where *N*_*pre*_ is the number of presynaptic neurons (PYRs or INs), *N*_*post*_ is the number of postsynaptic neurons (PYRs or INs) and *mean connectivity* is a parameter that is randomly varied between simulations. Please note that, due to the normalization, the fact that the two original distributions have a different mean is irrelevant. This process was repeated until the simulated number of connections had no negative values and no values exceeded the maximum amount of possible synaptic partners (*N*_*pre*_ or *N*_*post*_). In the case of uncorrelated incoming and outgoing number of synapses, the same procedure was then repeated for the distribution of pre/post-synaptic partners. In simulations in which the number of incoming and outgoing synapses was correlated, the number of presynaptic connections was drawn using a random number generator (numpy function *rng*.*choice*), in which the maximum value (a, following the nomenclature of *rng*.*choice*) was set equal to the number of pre/post-synaptic neurons, the number of values to draw was equal to *tot. connections* (size, following the nomenclature of *rng*.*choice*) and the probabilities associated with each presynaptic neuron were equal to the number of outgoing connections of that neuron (p, following the nomenclature of *rng*.*choice*). This resulted in a Pearson correlation of the number of incoming and outgoing synapses ∼0.85, in line with what has been described in Drosophila^92^. Finally, to generate a connectivity matrix in which each neuron had the desired amount of incoming and outgoing connections, we used the *directed_havel_hakimi_graph* function from the *networkx* python package.

#### Size of synapses

The distribution of synaptic sizes was simulated according to a normal (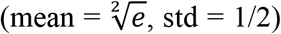, std = 1/2) or log-normal (mean of the underlying normal = 0, std of the underlying normal = 1) distribution. Please note that it can be analytically shown that the two distributions have the same mean. These distributions were then scaled by different scalars (see table 1) according to the specifics of the connected populations.

#### Synaptic plasticity

For simulations with synaptic plasticity, synaptic parameters were set to their extreme configuration (lognormal distribution of synaptic weights and number, and correlated amount of incoming and outgoing number of synapses). A set of other parameters controlling the network architecture was systematically varied across a biologically-constrained range, analogously to simulations without synaptic plasticity (see *Random variables*). In these networks, we first simulated 10s with frozen synaptic weights, and 40s with synaptic plasticity.

PYR-PYR excitatory synapses were plastic according to a classic asymmetric Hebbian plasticity rule^57^ that can be summarized as follows:

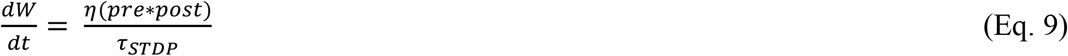

Where *W* is a PYR-PYR synaptic weight, *pre* and *post* are the pre- and postsynaptic activity, *η* is the learning rate, and *τ*_*STDP*_ is the decay time constant of the plasticity rule.

In practice, synaptic weights from a pre-synaptic neuron i to a post-synaptic neuron j (*W*_*ij*_) were updated at every pre- and post-synaptic event occurring at time *t*_*i*_ and *t*_*j*_ such that:

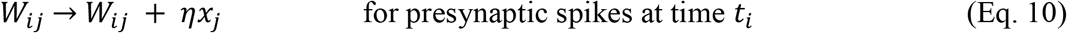

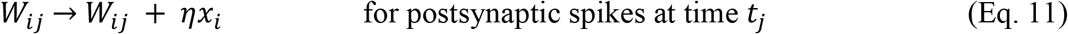

Where *x*_*i*_ and *x*_*j*_ are the pre- and post-synaptic trace.

*x*_*i*_ was updated with each spike *x*_*j*_ → *x*_*j*_+ 0.01 and decayed according to:

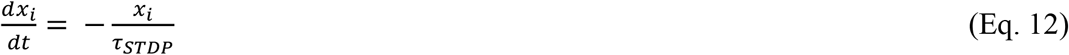

*x*_*j*_ was updated with each spike *x*_*j*_ → *x*_*j*_ − 0.01 and decayed according to:

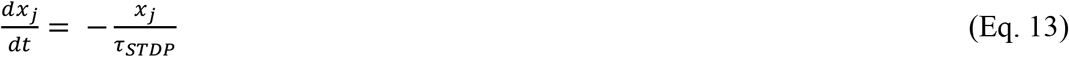

Synaptic weights were clipped within the 0-10 range.

IN-PYR inhibitory synapses were plastic according to a symmetric plasticity rule as previously described^56^. This rule can be summarized as follows:

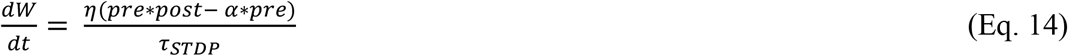

Where *W* is an IN-PYR synaptic weight, *pre* and *post* are the pre- and postsynaptic activity, *α* is the target rate for the postsynaptic PYR, drawn from a log-normal distribution, *η* is the learning rate, and *τ*_*STDP*_ is the decay time constant of the plasticity rule.

Synaptic weights *W*_*ij*_ were updated at every pre- and post-synaptic event occurring at time *t*_*j*_ and *t*_*i*_ such that:

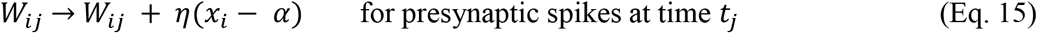

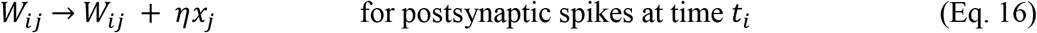

Where *x*_*i*_ and *x*_*j*_ are the pre- and post-synaptic trace that increased with each spike x*x x* → *xx* + 1 and decayed according to:

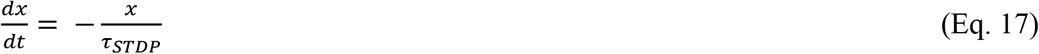

Synaptic weights were clipped within the 0-100 range.

#### Simulated SUA statistics and quality of model fit

For each network, we extracted the simulated spike times and computed the same 7 parameters as for the experimental data (described in *SUA firing statistics and shape distribution parameters*): skewness, kurtosis and Gini coefficient of firing rate and STTC, and the correlation between the log-transformed firing rate and STTC. To compute the quality of the model fit, we extracted the median of these 7 parameters from the P4-12 PFC dataset. For each simulation, we then computed the Euclidian distance (MATLAB function *pdist*) between the median of the experimental data and the 7D coordinates of that specific simulation. To ensure that each metric contributed equally to the distance from the experimental data, kurtosis and skewness of firing rate and STTC were divided by their maximum value. This normalized them to a range comprised between 0 and 1, as the Gini coefficient.

### Statistical modeling

Statistical modeling was carried out in the R environment. All the scripts and the processed data on which the analysis is based will be available on the following github repository: https://github.com/mchini/Chini_et_al_Preconfigured.

Nested data (figures 1, 5 and 7) were analyzed with (generalized) linear mixed-effects models (*lmer* and *glmer* functions of the *lme4* R package^101^) with “mouse” as random effect. Non-nested data were analyzed with linear models (*lm* function). Regression on data that was better fit by an exponential curve (figure 1) was carried out with a generalized linear mixed-effect models with the following parameters: family=Gamma, link=log. For ease of interpretability and consistency with the other distribution shape parameters, kurtosis of firing rate and STTC, which followed an approximately log-normal distribution, was instead log-transformed and analyzed with a linear model.

Statistical significance for linear mixed-effects models was computed with the *lmerTest* R package^102^ and the *summary* (type III sums of squares) R function. Statistical significance for linear models was computed with the *summary* R function.

When possible, model selection was performed according to experimental design. When this was not possible, models were compared using the *compare_performance* function of the *performance* R package^103^, and model choice was based on an holistic comparison of AIC, BIC, RMSE and R2.

Model output was plotted with the *plot_model* (type=‘pred’) function of the *sjPlot* R package^104^. 95% confidence intervals were computed using the *confint* R function. Post hoc analysis was carried out using the *emmeans* and *emtrends* functions of the *emmeans* R package^105^.

